# Anatomical and functional mapping of vagal nociceptive sensory nerve subsets innervating the mouse lower airways by intersectional genetics

**DOI:** 10.1101/2025.05.18.654723

**Authors:** Mayur J Patil, Justin Shane Hooper, Seol-Hee Kim, Parmvir K Bahia, Nika Pavelkova, Sanjay S Nair, Teresa S Darcey, Karina V Lurye, Meghana Madaram, Jailene Fiallo, Stephen H Hadley, Thomas E Taylor-Clark

## Abstract

Most vagal sensory afferents innervating the lower airways are activated by noxious stimuli including irritants (e.g. TRPV1 agonist capsaicin) and inflammatory mediators, causing nociceptive cardiorespiratory reflexes (e.g. cough, bronchospasm, changes in respiratory drive and heart rate). Vagal ganglia are comprised of embryologically distinct nodose and jugular neurons, but little is known of their specific contribution to nociceptive reflexes. Using a novel TRPV1^Flp^ mouse in combination with P2X2^Cre^, Tac1^Cre^, intersectional reporter mice and AAV we mapped and modulated distinct nociceptive afferents. TRPV1^+^P2X2^+^ neurons were found exclusively in the nodose ganglion and were activated by αβmATP and capsaicin but rarely expressed Tac1. TRPV1^+^P2X2^+^ fibers innervated the lungs (many projected into the alveoli) but not the trachea. Centrally they innervated the nucleus tractus solitarius (nTS). >90% of TRPV1^+^Tac1^+^ neurons were found in the jugular ganglion and were activated by capsaicin but not αβmATP. TRPV1^+^Tac1^+^ fibers innervated the lungs (although none projected into the alveoli) and the trachea submucosa. They terminated solely in the paratrigeminal complex (Pa5). Many TRPV1^−^Tac1^+^ neurons were found in both nodose and jugular ganglia that innervated the trachea and large pulmonary airways. These projected to both nTS and Pa5. Using intersectional chemogenetics we selectively stimulated lower airway afferent subsets using intravenous injections of clozapine-N-oxide (CNO). Activation of TRPV1^+^, TRPV1^+^P2X2^+^ or TRPV1^+^Tac1^+^ fibers evoked bradycardia and bradypnea. Activation of Tac1^+^ fibers evoked tachycardia and tachypnea. Activation of vagal TRPV1^−^Tac1^+^ neurons only evoked tachycardia. These data show the distinct innervation patterns and reflex function of multiple nociceptive vagal afferent subsets.

## Introduction

Sensory afferent nerves provide real-time information to the organism of its internal and external environment in order to initiate reflexes and behaviors that ultimately serve to maintain physiological homeostasis ^1^. Given that the environment comprises of diverse stimuli that trigger a variety of responses it is not surprising that sensory afferents are heterogeneous with respect to gene expression and neuroanatomy ^2,3^. The vagus nerve provides sensory innervation to almost all visceral organs and mediates many reflexes that regulate respiratory, cardiovascular, gastrointestinal, genitourinary, metabolic and inflammatory systems ^4–6^. Most vagal afferents (>80%) are unmyelinated, slowly-conducting C fibers which are largely quiescent under normal conditions, but instead are activated by noxious or potential noxious stimuli ^7^. These so-called nociceptive afferents express numerous receptors that are activated by irritants (e.g. capsaicin-sensitive Transient Receptor Potential (TRP) Vanilloid 1 (V1)), inflammatory mediators and other noxious chemical and physical conditions ^3^. The remaining vagal afferents are myelinated, fast-conducting A fibers, many of which are low-threshold mechanoreceptors that are typically insensitive to noxious stimuli.

The lower airways (trachea, bronchi and lungs) receive >90% of their sensory innervation from the vagal ganglia ^8–10^. Activation of nociceptive afferents within the lower airways (e.g. via the administration of capsaicin, the pungent ingredient in chili peppers) triggers reflexes and symptoms that mimic infectious or inflammatory airway diseases: cough, dyspnea, hypersecretion, bronchospasm, changes in respiratory drive, heart rate and blood pressure ^11,12^. Furthermore, airway nociceptive afferents appear to be essential for airway hyperreactivity ^13–16^ and pulmonary dysfunction in pneumonia ^17^. Nevertheless, there is surprising heterogeneity in airway nociceptive reflexes ^18–22^, suggesting that multiple functionally distinct nociceptive subsets innervate the lower airways.

Importantly, vagal sensory neurons are derived from two distinct embryological sources: placodes-derived neurons in the nodose ganglion, and neural crest-derived neurons in the jugular ganglion ^23^. Both ganglia contain neurons that are likely to be nociceptive (i.e. they express TRPV1, the wasabi-sensitive TRP ankyrin 1 (A1), and the bradykinin B2 receptor) and neurons likely to be non-nociceptive. Nevertheless, transcriptomics studies have identified substantial phenotypic differences between nodose and jugular neurons ^3,24,25^: for example, nodose neurons express the master regulator Phox2b, the neurotrophin TrkB receptor and autacoid receptors P2X2 and 5HT3; whereas jugular neurons express Prdm12 and the neurotrophin TrkA receptor but do not express P2X2 or 5HT3. Tac1, the gene that encodes the neuropeptide substance P, is largely expressed in the jugular ganglia – often in TRPV1^+^ neurons ^7^. However, Tac1 is also expressed in some nodose TRPV1^−^ neurons.

Previous studies have used cre/lox approaches to map vagal afferents. Fluorescent reporters expressed in afferent subsets under the control of Phox2b^Cre^, TRPV1^Cre^, P2X2^Cre^ and Tac1^Cre^ suggest differences in innervation patterns within the lower airways and the medulla, which may provide a neuroanatomical explanation for the heterogeneity of reflex responses evoked from the airways. In the lung, Phox2b^+^, P2X2^+^ and TRPV1^+^ fibers innervate conducting airways, alveoli and pulmonary blood vessels, whereas Tac1^+^ fibers innervate only conducting airways and blood vessels ^9,10,26^. In the medulla, Phox2b^+^ and P2X2^+^ fibers terminate in the nucleus tractus solitarius (nTS), whereas TRPV1^+^ and Tac1^+^ fibers terminate in both the nTS and the paratrigeminal complex (Pa5) ^7,27,28^. Nevertheless, it is important to note that no single gene can be used to identify a discrete vagal afferent subset thus the link between genetically identified subsets and their function has yet to be established.

Here, we have used an intersectional genetics approach to anatomically and functionally map vagal nociceptive afferents. Combining a novel TRPV1^Flp^ knockin mouse strain with either P2X2^Cre^ or Tac1^Cre^ allows for the identification and manipulation of afferent subsets identified by the intersection of two genes: e.g. TRPV1^+^P2X2^+^ fibers (nodose TRPV1^+^ nociceptors) and TRPV1^+^Tac1^+^ (jugular TRPV1^+^ nociceptors).

## Methods

### Animals and genotyping

The gene for the murine TRPV1 receptor (NCBI Reference Sequence: NM_1001445.2) is located on chromosome 11. Sixteen exons have been identified, with the TGA stop codon in exon 16. To develop a knock-in mouse that expresses the codon-optimized Flp_O_ recombinase dependent on TRPV1 expression, the TGA stop codon was replaced with a 2A-Flp cassette (Fig. 1). The targeting vector homology arms were generated by high fidelity Taq PCR using BAC clone RP23-30N20 and RP23-152H19 from the C57BL/6J library as template. The targeting vector was assembled with recombination sites and selection markers: Neomycin resistance gene (Neo^R^) flanked by self-deletion anchor (SDA) sites for positive selection and diphtheria toxin A fragment gene (DTA) for negative selection. Correct targeting vector synthesis was confirmed by appropriate digestion by restriction enzymes. The linearized vector was subsequently delivered to C57BL/6 ES cells via electroporation, followed by drug selection, PCR screening, and Southern Blot confirmation. After gaining 94 neomycin-resistant clones, 14 potentially targeted clones were confirmed, 6 of which were expanded for Southern Blotting. After confirming correctly targeted ES clones via Southern Blotting, clones were selected for blastocyst microinjection, followed by founder production. Founders were confirmed as germline-transmitted via crossbreeding with wild-type C57BL/6J. All aspects of knock-in mouse development were performed by Cyagen US Inc (California). Founders were mated to produce heterozygous and homozygous *TRPV1^tm1.1(flp)Ttc^*(MGI:8161908) mice (*TRPV1^Flp^*) in expected Mendelian proportions. These mice express TRPV1-2A-Flp_O_ from the endogenous TRPV1 gene. Upon translation, the 2A peptide self-cleaves to release TRPV1 and Flp as separate peptides. *TRPV1^Flp^*mice develop normally and were observed to have no apparent phenotype. Homozygous *TRPV1^Flp^* were crossed with Flp-sensitive *R26^ai65f^* reporter mice (B6.Cg-*Gt(ROSA)26Sor^tm65.2(CAG–tdTomato)Hze^*/J, #032864, Jackson Laboratory) ^29^ that possess an FRT-stop-FRT-tdTomato sequence in the ROSA26 locus, to produce heterozygous *TRPV1^Flp/+^R26^ai65f/+^* mice, which express tdTomato in Flp-expressing cells. Homozygous *TRPV1^Flp^* were also crossed with Flp-sensitive *R26^FLTG^* reporter mice (B6.Cg-*Gt(ROSA)26Sor^tm1.3^*(CAG–tdTomato,–EGFP)*^Pjen^*/J, #026932, Jackson Laboratory) to produce heterozygous *TRPV1^Flp/+^R26^FLTG/+^*mice. The ROSA26 locus of *R26^FLTG^* mice has an FRT-stop-FRT-LoxP-tdTomato-stop-LoxP-GFP sequence, which expresses tdTomato in Flp^+^Cre^−^ cells and GFP in Flp^+^Cre^+^ cells. In some cases *TRPV1^Flp/+^R26^FLTG/+^*mice were back-crossed to produce homozygous mice for one or both alleles. *TRPV1^Flp^*were crossed with *P2X2^Cre^* mice (*P2rx2^tm1.1^*(cre)*^Ttc^*) ^27^ to produce *TRPV1^Flp/+^P2X2^Cre/+^* mice, which were then back-crossed to homozygosity. Homozygous *TRPV1^Flp^P2X2^Cre^* mice were crossed with homozygous *R26^FLTG^* mice to produce heterozygous *TRPV1^Flp/+^P2X2^Cre/+^R26^FLTG/+^* (*TRPV1^Flp^P2X2^Cre^-FLTG*) mice. *TRPV1^Flp^* were crossed with *Tac1^Cre^* mice (B6.129S-*Tac1^tm1.1(cre)Hze^*/J, #021877, Jackson Laboratory) ^30^ to produce *TRPV1^Flp/+^Tac1^Cre/+^* mice, which were then back-crossed to homozygosity. Homozygous *TRPV1^Flp^Tac1^Cre^*mice were crossed with homozygous *R26^FLTG^* mice to produce heterozygous *TRPV1^Flp/+^Tac1^Cre/+^R26^FLTG/+^* (*TRPV1^Flp^Tac1^Cre^-FLTG*) mice. We also crossed the *P2X2^Cre^* mice and *Tac1^Cre^*mice with transgenic CAG-loxP-STOP-loxP-Gq-DREADD mice (*B6N;129-Tg(CAG-CHRM3*,-mCitrine)1Ute*/J, #026220, Jackson Laboratory, formally known as *Gt(ROSA)26Sor^tm2(CAG-CHRM3*,-mCitrine)Ute^/J*) to produce heterozygous *P2X2^Cre^-Dq* and *Tac1^Cre^-Dq* mice that express the clozapine-N-oxide (CNO)-sensitive ‘activating’ DREADD hM3Dq receptor in P2X2- and Tac1-expressing cells, respectively. Homozygous knock-in *Pirt^Cre^* mice (*Pirt^tm3.1(cre)Xzd^*, kind gift from Dr. Xinzhong Dong, Johns Hopkins University) ^31^ and homozygous knockin *TRPV1^Cre^* mice (*B6.129X1-Trpv1^tm1(cre)Bbm^*/J, #017769, Jackson Laboratory) ^32^ were used for intraganglionic injections. Lastly, C57BL/6J mice (#000664, Jackson Laboratory) were used as control mice for the chemogenetic studies.

**Figure 1:**
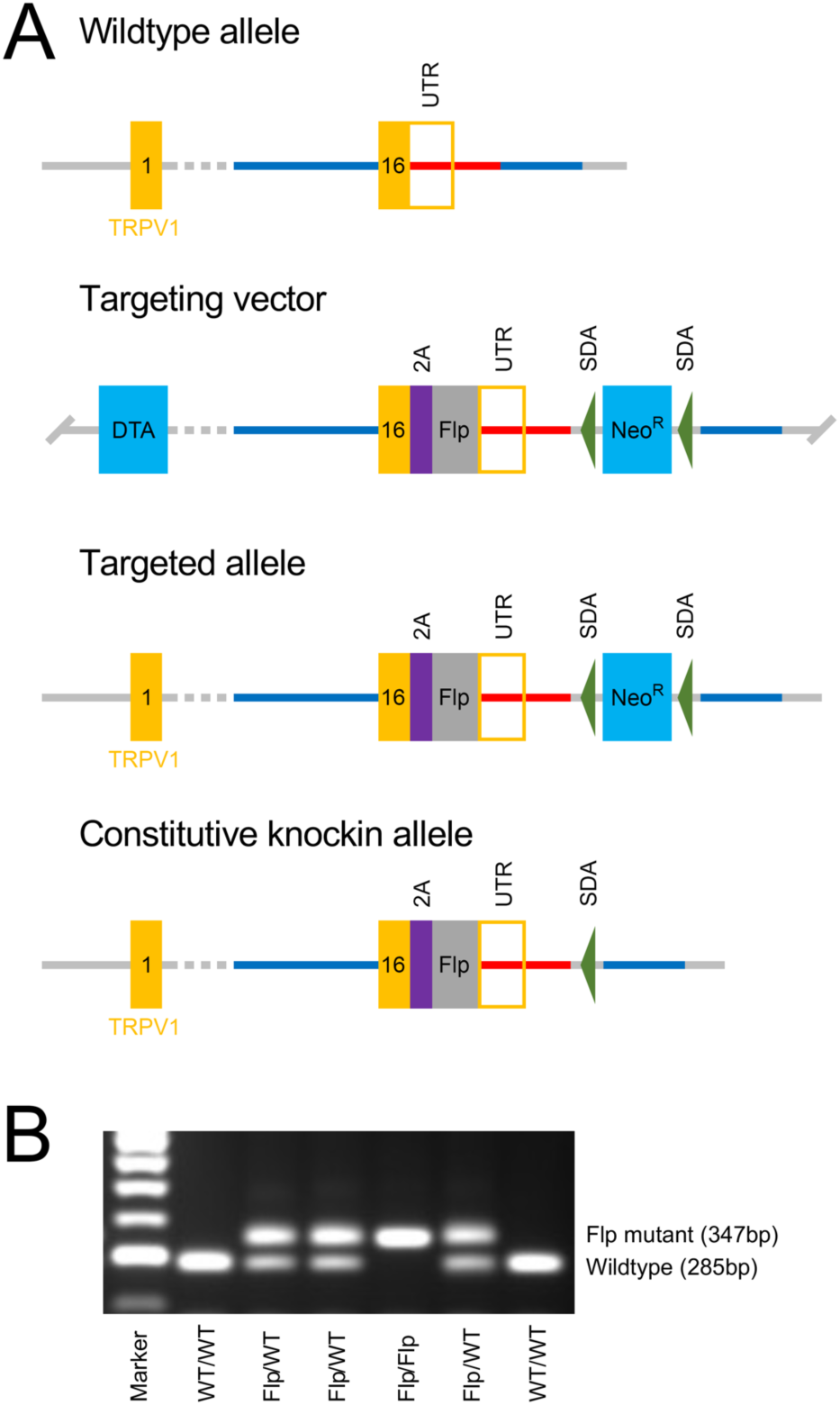
Development of the knock-in *TRPV1^Flp^* mouse. A, targeting strategy for the replacement of the TRPV1 TGA stop codon with a 2A-Flp_O_ cassette. Homology arms (blue and red lines) were generated for the TRPV1 gene (exons and 3’ untranslated region (UTR) in orange). Homology arms of targeting vector include a neomycin resistance gene (Neo^R^) flanked by self-deletion anchor (SDA) sites for positive selection. A Diphtheria toxin A fragment gene (DTA) was placed in a non-homologous region of targeting vector as a negative selection for non-homologous recombination. B, PCR of *trpv1* gene in offspring from a pairing of heterozygous *TRPV1^Flp^* mice. As expected, offspring have a mendelian distribution of mutant (i.e. *TRPV1^Flp^*, at 347 bp) and wildtype (at 285 bp) alleles.

Both male and female mice (6 to 8-week-old) were used for experiments. Offspring were weaned at 21 postnatal days, and up to 4 littermates were housed per cage under normal condition (20 ⁰C, a 12 hour dark/light cycle). Mice were provided with standard rodent chow and water *ad libitum*. All procedures were in accordance with the animal protocols approved by the University of South Florida Institutional Animal Care and Use Committee.

### Intraganglionic injection with adeno-associated virus (AAV)

Recombinase-expressing mice (all homozygous) were administered intraganglionic injections of AAV9 into either the vagal ganglia or the dorsal root ganglia (DRG) to drive reporter (unilateral injection) or DREADD hM3Dq receptor (bilateral injections) expression. AAV used in this study (Table 1) had recombinase sensitive sites within their DNA sequence to facilitate cell-type specific expression ^33^. Motifs that are recombined to switch nonsense sequences to sense sequences under the control of Cre recombinase and Flp recombinase are termed “Con” and “Fon”, respectively. Similarly, motifs that are recombined to switch sense sequences to nonsense sequences under the control of Cre recombinase and Flp recombinase are termed -Coff- and - Foff-, respectively. As previously described, the targeted insertion of introns into the vector sequence allows for sequential recombination of a given gene by Cre and Flp ^33^: “ConFon”, “ConFoff” and “CoffFon” intersectional motifs.

**Table 1.** AAVs used in this study.

| Virus | Titer | Source | Cat# |
| --- | --- | --- | --- |
| AAV9-Con-tdTomato | $2.1 \times 10^{13}$ GC/mL | Addgene | 28306 |
| AAV9-Con-EGFP | $2.5 \times 10^{13}$ GC/mL | Addgene | 510502 |
| AAV9-ConFon-mCherry | $4.6 \times 10^{13}$ GC/mL | Princeton | VC362 |
| AAV9-ConFoff-mCherry | $6.4 \times 10^{13}$ GC/mL | Princeton | VC366 |
| AAV9-CoffFon-mCherry | $1.0 \times 10^{14}$ GC/mL | Princeton | VC367 |
| AAV9-Con-hM3Dq-mCherry | $2.3 \times 10^{13}$ GC/mL | Addgene | 44361 |
| AAV9-ConFon-hM3Dq-mCherry | $1.8 \times 10^{13}$ GC/mL | Princeton | VC363 |
| AAV9-ConFoff-hM3Dq-mCherry | $3.1 \times 10^{13}$ GC/mL | Princeton | VC368 |

### Vagal ganglia microinjection

Using a procedure that has been described elsewhere ^10^, mice were anesthetized with ketamine (50mg/kg) and dexmedetomidine (0.5 mg/kg) via intraperitoneal injection. Approximately 2 cm of incision was made over a shaved superficial portion of the masseter muscle area. The skin was retracted, and the vagus nerve was located and then separated from the common carotid artery and the anterior laryngeal nerve using a cotton tip. The virus microinjection assembly consisted of a pulled glass micropipette (∼20μm tip diameter) attached to a 1mL syringe via plastic tubing. The tip of the micropipette was gently inserted into the exposed vagal ganglia and then 0.7μL of AAV9 was injected using ∼0.5 p.s.i. pressure. Three to 6 weeks later the mice were used for analysis.

### Dorsal root ganglia microinjection

The thoracic (T1-T3) DRG were unilaterally injected using a procedure described elsewhere ^10^. Mice were anesthetized with ketamine (50mg/kg) and dexmedetomidine (0.5 mg/kg) via intraperitoneal injection. The neck area was shaved, and a 4 cm incision was made to expose the dorsal cervical region. The last cervical vertebra was identified and used as a guide. The underlying muscle fibers were separated to expose the T1, T2, and T3 intervertebral space. The spinal nerves were identified and followed to the beginning of the ganglia. A pulled glass micropipette (∼20μm tip diameter) was prefilled with 0.75 μl of AAV9 virus then injected into each DRG using ∼0.5 p.s.i. pressure. Three to six weeks later, mice were used for analysis.

### Tissue collection and immunohistochemistry

Mice were euthanized by CO_2_ inhalation and transcardially perfused with ice-cold PBS. Vagal ganglia and DRG from T1 to T4 were dissected out and fixed for 30 mins in 3.7% formaldehyde (FA) at 4⁰C. Brainstem were dissected out and fixed for 4 hours in 3.7% FA at 4 ⁰C.

All tissues to be sliced using a cryostat were first cryoprotected in sucrose (20% for ganglia, 30% for brainstem). Ganglia were sectioned sagitally in 20μm slices, and brainstem were sectioned coronally in 40μm slices, collected onto Superfrost plus slides. Slides were air-dried at room temperature in the dark overnight. Slides were washed with PBS three times for 10 mins, and tissue was permeabilized with 0.3% Triton X-100 in PBS (PBSTx) for 15 mins followed by blocking for 1 hour with 1% bovine serum albumin (BSA)/10% Donkey Serum (DS)/0.3% PBSTx. Tissue was incubated with primary antibodies (Table 2) diluted in blocking buffer overnight at 4 ⁰C. After washing with 0.2% Tween 20 in PBS (PBST) three times for 10 mins, the tissue was incubated with secondary antibodies (Table 2) in 1% BSA/5% DS in 0.2% PBST for 1 hour. The tissue was washed with 0.2% PBST three times for 10 mins and then, in some cases, counterstained with NeuroTrace^TM^ fluorescent Nissl Stain for 1 hour at 1:300 dilution in PBS. After washing with PBS, slides were air-dried and mounted with DPX mounting medium (Sigma).

**Table 2.** Antibodies used in this study.

| **Antibodies** | **Host** | **Dilution** | **Catalog No.** | **Source** | **RRID** |
| --- | --- | --- | --- | --- | --- |
| Anti-DsRed | Rabbit | 1:500 | 632496 | Takara Bio | AB_10013483 |
| Anti-RFP | Rabbit | 1:1:200 | 600-401-379 | Rockland | AB_2209751 |
| Anti-GFP | Chicken | 1:1000 | ab13970 | Abcam | AB_300798 |
| Anti-GFP | Chicken | 1:200 | NB100-1614 | Novus | AB_10001614 |
| Anti-E-cadherin | Rat | 1:300 | ab11512 | Abcam | AB_298118 |
| Anti-α-smooth muscle actin | Goat | 1:300 | SAB2500963 | MilliporeSigma | AB_10603763 |
| Anti-CGRP | Rabbit | 1:300 | C8198 | MilliporeSigma | AB_259091 |
| Neurotrace™<br>435/455 | N/A | 1:300 | N21479 | Invitrogen | AB_2572212 |
| Neurotrace™<br>500/525 | N/A | 1:300 | N21480 | Invitrogen | SCR_013318 |
| Alexa fluor 488<br>anti-chicken<br>immunoglobulin | Goat | 1:500 | A11039 | Invitrogen | AB_2534096 |
| Alexa fluor 488<br>anti-rabbit<br>immunoglobulin | Donkey | 1:500 | A21206 | Invitrogen | AB_2535792 |
| Alexa fluor 546<br>anti-rabbit<br>immunoglobulin | Donkey | 1:500 | A10040 | Invitrogen | AB_2534016 |
| Alexa fluor 546<br>anti-rat<br>immunoglobulin | Goat | 1:500 | A11081 | Invitrogen | AB_141738 |
| Alexa fluor 647<br>anti-chicken<br>immunoglobulin | Goat | 1:500 | Ab150171 | Abcam | AB_2921318 |
| Alexa fluor 647<br>anti-rat<br>immunoglobulin | Goat | 1:500 | A21247 | Invitrogen | AB_141778 |
| Alexa fluor 647<br>anti-goat<br>immunoglobulin | Chicken | 1:500 | A21469 | Invitrogen | AB_2535872 |
| Alexa fluor 647<br>anti-rabbit<br>immunoglobulin | Donkey | 1:500 | A31573 | Invitrogen | AB_2536183 |
| Biotin-XX<br>conjugate anti-rabbit<br>immunoglobulin | Goat | 1:100 | B2770 | Molecular<br>Probes | AB_2536431 |

Complete lungs with attached bronchi and trachea were dissected out and fixed in 3.7% FA overnight at 4 ⁰C with gentle agitation. Lungs were washed with ice-cold PBS three times for 30 mins, followed by cryoprotection in 30% sucrose solution. The lungs were flushed with PBS three times and inflated with 2% low melting agarose solution. After the agarose solution had solidified, the lung lobes were separated and snap frozen with OCT compound for cryosection. 80μm lung slices were collected in cryoprotectant-filled wells and stored in -20 ⁰C. Using identical blocking solutions, permeabilizing solutions and antibody solutions (see above), lung slices were washed in PBS three times for 10 mins and permeabilized for 20 mins. Lung slices were then blocked for 1.5 hours followed by primary antibody incubation overnight at 4 ⁰C. The slices were then washed three times for 20 mins and incubated with secondary antibodies for 2 hours. The slices were washed again and mounted onto glass slides with VECTASHIELD^®^ Antifade Mounting Medium with DAPI.

The trachea were dissected out and cut longitudinally along the ventral side. The tracheal ends were pinned in a Sylgard covered dish and fixed in 3.7% FA overnight at 4 ⁰C. Whole tissue tracheal staining was performed in 2 ml tubes. The trachea were washed in filtered phosphate buffered saline (PBS) between staining steps, which was performed in 15-ml tubes on the rotor at room temperature. The trachea were permeabilized in 1% Tween 20 (Sigma-Aldrich, St. Louis, MO) in filtered PBS at room temperature for 6 h, washed in filtered PBS (3 × 20 min on rotator). The trachea were blocked with BSA/PBS individually at 4°C for 4 hours. The trachea was washed 5 × 20 min on the rotator and then incubated with primary antibodies diluted in BSA/PBS for 48h at 4°C. The trachea was then washed in filtered PBS (10 × 30 min) and incubated with secondary antibodies in BSA/PBS overnight at room temperature, then washed in filtered PBS (3 × 20 min), incubated in anti-fade (pH 8.6) glycerol (Sigma-Aldrich) for 24h at room temperature, and stored at 4 °C. The trachea were mounted on superfrost coated slides (FisherScientific) with the lumen surface facing up and pressed onto the slide with 1 pound weight overnight before imaging.

Images were taken with Andor Dragonfly spinning disk confocal microscope equipped with a Zyla 4.2 PLUS sCMOS camera (2048 x 2048 pixel with 6.5μm pixel size). The pinhole size was 25μm. We used either a 10x UPLSAPO (0.4NA), a 20x UPLSAPO (0.75NA) or a 40x UPLSAPO (1.25NA, silicone oil immersion) objective was used, depending on the study. Fluorophores were excited by laser wavelengths at 405nm, 488nm, 561nm or 637nm. Z-stacked multi-tile images were stitched using either Fusion software or Imaris Stitcher. All 3D images were further processed using Imaris software. The identification of anatomical structures and subnuclei were based on the mouse brain map ^34^. In mice the nodose vagal ganglion and the jugular vagal ganglion are partially fused into a single structure. Nevertheless, these two ganglia can be approximately discriminated visually in sagittal slices of the vagal ganglia, as shown previously ^7,27,35^. Cell counts and somal diameter of neurons within the vagal ganglia and DRG were obtained using Fiji software. The somal areas of vagal and DRG afferent subpopulations were compared using unpaired t tests.

### Sensory ganglia dissociation

Mice were killed by CO_2_ asphyxiation followed by exsanguination. DRG or vagal ganglia were isolated in Ca^2+^-free, Mg^2+^-free Hank’s buffered saline solution HBSS, then incubated in HBSS containing type 1 collagenase (2mg/ml) and dispase II (2mg/ml), then mechanically dissociated with fire-polished pipettes. Individual neurons were washed and resuspended in L-15 media supplemented with 10% fetal bovine serum (FBS), 100 U/ml penicillin and 100 µg/ml streptomycin then plated onto poly-D-lysine and laminin coated coverslips. Neurons were incubated at 37°C in antibiotic-free L-15 media supplemented with 10% FBS and used within 24 hours.

### Single neuron RT-PCR

Coverslips with dissociated vagal sensory neurons were superfused with Krebs bicarbonate solution (KBS, in mM: 118 NaCl, 5.4 KCl, 1 NaH_2_PO_4_, 1.2 MgSO_4_, 1.9 CaCl_2_, 25 NaHCO_3_, and 11 dextrose, gassed with 95% O_2_-5% CO_2_). Reporter-expressing neurons were identified by fluorescence (GFP – 475 nm excitation, 520 nm emission; tdTomato – 554 nm excitation, 581 nm emission) and an Olympus BX51WI microscope. Individual neurons were collected with a borosilicate glass pipette by applying a gentle negative pressure. Each single neuron was stored in PCR tube containing 1μl RNase OUT and stored at -80°C. The KBS surrounding the vagal neurons was sampled (∼2μl) as a negative control for the RT-PCR.

First-strand cDNA was synthesized with SuperScript®III First-Strand Synthesis System for RT-PCR by following the manufacturer’s instructions. Primers and dNTP mix were added into each neuron-containing PCR tube. Samples were incubated at 75°C for 10min and then placed on ice for at least 1 min. cDNA synthesis mix containing 10x RT buffer, MgCl_2_, DTT and SuperScript III RT was added into each sample. Samples were incubated at 50°C for 50min, followed by 85°C for 5min. cDNA was stored at -20°C until used for PCR amplification. cDNA was amplified with HotStarTaq DNA polymerase for 50 cycles of denaturation at 94°C for 30s, followed by annealing at 60°C for 30s and extension at 72°C for 1min. Customized intron-spanning primers for mouse TRPV1, P2X_2_, preprotachykinin A (PPT-A) and vesicular glutamate transporter 2 (VGLUT2) (Table 3) were used as previously reported ^35,36^. Products were visualized in 1.5% agarose gel with GelRed, with a 100-bp marker. VGLUT2 was used as a positive control.

**Table 3:**
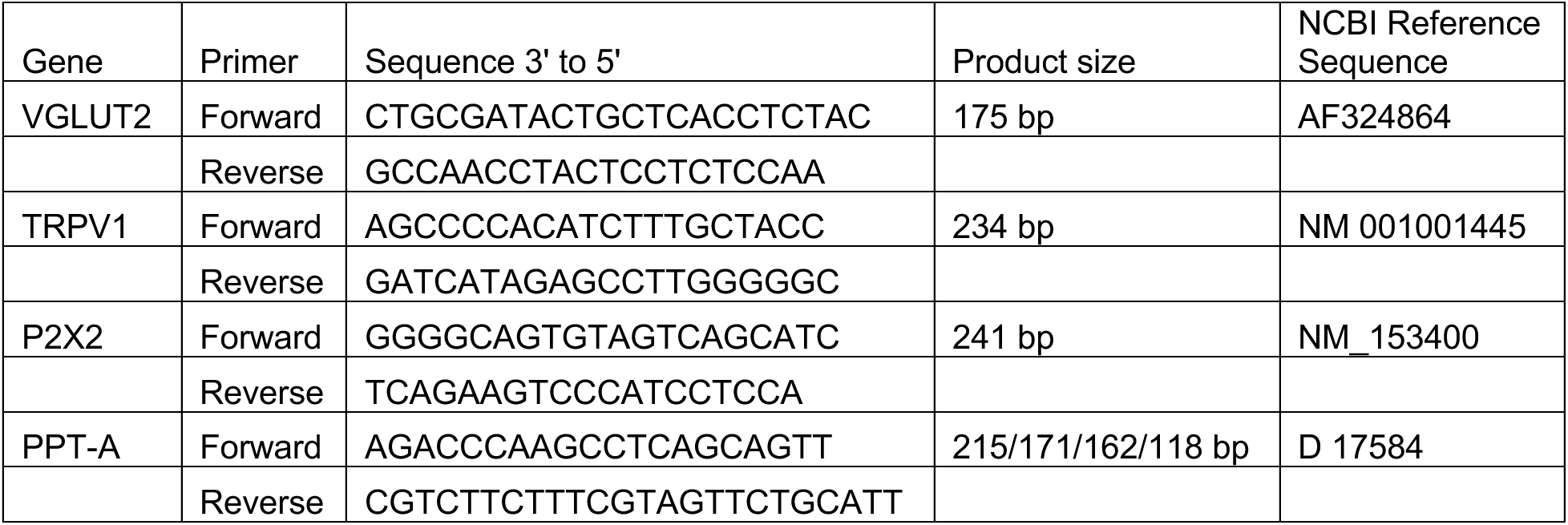
Primers used for single cell reverse transcription polymerase chain reaction in mouse neurons.

### FURA-2AM Ca^2+^ imaging and analysis

Neuron-coated coverslips were incubated with 4μM FURA-2AM for 30-60 minutes at 37°C. Coverslips were loaded into a chamber on a Nikon Eclipse Ti inverted microscope, fitted with a CFI Super Fluor 10x objective and a CoolSnap HQ2 camera (Photometrics), and perfused with heated (33-34°C) HEPES buffer (154mM NaCl, 1.2mM KCl, 1.2mM MgCl_2_, 2.5mM CaCl_2_, 5.6mM D-Glucose). Cells were subjected to fluorescent light delivered by a 300W (5000 lumens) PE300BFA Xenon parabolic bulb housed in a Lambda LS, with appropriate excitation and emission filters. Brightfield and fluorescence images were collected to facilitate subsequent analysis and the determination of GFP (470nm excitation, 525nm emission) and tdTomato expression (535nm excitation, 610nm emission), respectively. Changes in [Ca^2+^]_i_ were monitored every 6 seconds using sequential excitation at 340nm and 380nm (510nm emission) and evaluated ratiometrically using the 340/380 ratio (R). All drugs were diluted in HEPES buffer. α,β methylene ATP (αβmATP, 10μM) was used to determine the functional expression of P2X2 ^27^, and capsaicin (1μM) was used to determine the functional expression of TRPV1 ^37^. Neurons were further characterized by response to KCl (75mM) prior to ionomycin (5μM), which evoked a maximal Ca^2+^ response. Region of interest (ROI) analysis for individual neurons was performed using Nikon Elements (Nikon, Melville, NY). ROIs with an unstable, high, or noisy baseline were eliminated from analysis. Neurons which failed to exhibit an increase in [Ca^2+^]_i_ to either capsaicin or KCl challenges (> 30% the ionomycin maximal response) were eliminated. An individual neuron was considered to be sensitive to a given agent if R_agent_ > (R_0_ + 2*SD_0_) + 0.015; where R_agent_ is the average 340/380 ratio during treatment, R_0_ is the average 340/380 ratio over 60s prior to treatment, and SD_0_ is the standard deviation of 340/380 ratios over 60s prior to treatment. Neurons were grouped by GFP and tdTomato expression and sensitivity to αβmATP and capsaicin. For each neuron, each time point responses was normalized by subtraction of the baseline (R_1_-R_0_). Normalized responses to a given stimulus were compared between different neuronal groups using unpaired t tests.

### Physiological measurements in anesthetized mice

Mice were anesthetized with ketamine (100mg/kg) and dexmedetomidine (0.5 mg/kg) via intraperitoneal injection, and the level of anesthesia was confirmed by lack of response to inter-digital pinch every 15 minutes. Puralube Vet Ointment (Dechra) was applied to the eyes to prevent irritation caused by desiccation before the mice were placed in a supine position on a temperature-controlled heating pad (Physiosuite, Kent Scientific). Core temperature was maintained at 36-38°C with the aid of a rectal thermometer. ECG was recorded using flexivent needle electrodes placed into the right pectoralis and left intercostal muscles. Similarly, dEMG was recorded using needle electrodes placed into the diaphragm approximately 1-cm apart. No incisions were made for electrode placement. If necessary, the electrodes were kept in place using medical tape. A 1-inch incision was made through the dermal layers on the ventral side of the mouse in line with the right sternomastoid beginning at the sternum (manubrium). The dermal layers were retracted, and surrounding tissues blunt dissected to expose the jugular vein. A butterfly IV (27G ½ inch, 24cm PU tubing) catheter was loaded with physiological saline (0.9% NaCl) and the needle was run through the pectoralis major and into the jugular vein. To minimize dead space, the butterfly was connected to the tubing with a PinPort (28G, Instech, PNP3M-F28). The line was held in place using tape after proper positioning of the i.v. was confirmed via flashing the line. If switching of the syringes was needed, the line was clamped with rubber hemostats before removing the syringe. Using the new syringe, the line was primed to remove any air from the line/coupler and then inserted before removing the hemostats for intravenous agonist administration (in 50 μL). The bolus given was equal to the calculated dead space in the tubing and all agonist boluses were immediately followed by a 50 μL flush of physiological saline (0.9% NaCl). The following protocols were used:

1) For C57BL/6J mice: after a 15-minute period, mice were administered PBS vehicle, then hM3Dq agonist clozapine-n-oxide dihydrochloride (CNO, 300μg/ml), then TRPV1 agonist capsaicin (3μg/ml), with the injections separated by 10 minutes.
2) For *P2X2^Cre^-hM3Dq* mice and *Tac1^Cre^-hM3Dq* mice: after a 15-minute period, mice were administered PBS vehicle, then CNO (3μg/ml), then capsaicin (3μg/ml), with the injections separated by 10 minutes.
3) For *TRPV1^Cre^* mice, *TRPV1^Flp^P2X2^Cre^* mice and *TRPV1^Flp^Tac1^Cre^*mice that have all been previously administered AAV9 vagal microinjections (hM3Dq-mCherry): after a 15-minute period, mice were administered PBS vehicle, then CNO (10μg/ml), then CNO (100μg/ml), then capsaicin (3μg/ml), with the injections separated by 10 minutes.

The ECG signal was amplified using a model P511 amplifier (Grass Instrumental Co., Quincy, Mass.) and filtered (band-pass 10 Hz to 10 kHz). The ECG trace was observed throughout the experiment via an oscilloscope and the signal was recorded continually, along with dEMG, on digital recorders (16-bit accuracy, 24- or 25kHz sampling frequency per channel) and later analyzed using the Spike 2 version 7 software (Cambridge Electronic Design, CED). A trigger threshold was set to distinguish R waves from P and T waves and the time between each R wave, R-R interval (RRi, in ms), was generated (such that the instantaneous heart rate per minute = 1000 * 60 / RRi). The dEMG signal was amplified using a model P511 amplifier and filtered (band-pass 10 Hz to 10 kHz). Noise derived from the cardiac cycle was subtracted from the dEMG signal using a CED custom script. The signal was then subjected to rectification, DC Remove (time constant of 0.004), RMS amplitude (time constant of 0.0004) and smoothing (time constant of 0.06 to 0.08). A trigger threshold was set to distinguish muscle contraction from background activity and the ECG artifact, and the interval between breaths (BBi, in ms) was generated (such that the instantaneous breathing rate per minute = 1000 * 60 / BBi). For each mouse, each RRi value was subtracted from the average RRi in the 20s prior to bolus injection to generate a ΔRRi value. The area under the curve (AUC) for the response on the ΔRRi against time graph was calculated for each stimulus, then divided by the duration of the stimulus response (in s). This yielded an overall ΔRRi for the entire response to a given stimulus for each animal. Similarly, for each mouse, each BBi value was subtracted from baseline to generate a ΔBBi value. The area under the curve (AUC) of the ΔBBi response against time was calculated, then divided by the response duration to yield an overall ΔBBi for the entire response to a given stimulus for each animal. ΔRRi and ΔBBi responses to CNO or capsaicin were compared to vehicle responses using paired T tests. ΔRRi and ΔBBi responses to vehicle and CNO in individual mice were visualized using rolling averages (averaged over 15 consecutive beats).

### qPCR validation of hM3Dq-mCherry expression

Following functional experimentation, vagal ganglia were dissected out and transferred into 700μl of Qiazol (Qiagen) along with 5 mm stainless steel beads (#69989, Qiagen) and immediately flash frozen in liquid nitrogen. The tissue was first disrupted in Qiazol using TissueLyser LT (#85600, Qiagen) for 5 minutes. The RNA was isolated using the RNeasy Plus mini kit (#74134, Qiagen), and the cDNA was synthesized using the SuperScript III RT (# 18080044, Invitrogen) with Oligo(dT)12-18 primer (#18418012, Invitrogen) and dNTP mix (#R72501). 70 ng of cDNA per reaction were used for RT-qPCR using FAM-labeled Taqman probes for GAPDH (Mm99999915_g1), VGLUT2 (Mm00499876 m1), and mCherry (Mr07319439_mr). The qPCR was performed in TaqMan gene expression master mix (#4369016, Life Technologies) for 45 cycles (95 °C for 1 s, 60 °C for 20 s). For recombinase-expressing mice that had received vagal injection of recombinase-sensitive AAV9 expressing hM3Dq-mCherry (n=40), the mean ± SEM Ct for GADPH was 28.4 ± 0.50, for VGLUT2 was 33.69 ± 0.96 and for mCherry was 37.87 ± 0.28 (representing an average ΔCt of 9.46 ± 0.45). No mCherry transcript was detected within 45 cycles in vagal ganglia taken from control mice (n=3). Thus the mCherry signal in each recombinase-expressing mice that had received vagal injection of recombinase-sensitive AAV9 expressing hM3Dq-mCherry was deemed sufficient for inclusion.

### Statistical analysis

All data was analyzed in GraphPad Prism 10. In all cases, a p value less than 0.05 was considered significant.

## Results

### Characterization of TRPV1^Flp^ reporter mouse

We generated a knock-in *TRPV1^Flp^* mouse (Fig. 1), which was initially crossed with the Flp-sensitive *R26^ai65f^* reporter strain to make *TRPV1^Flp/+^R26^ai65f/+^* mice, that were expected to express the red fluorescent tdTomato (tdT) in TRPV1-expressing cells. Frozen sections taken from the vagal ganglia and thoracic DRG showed selective tdT expression in 38.6% (1638 out of 4243) of vagal neurons and 30.0% (451 out of 1502) of DRG neurons (Fig. 2A, B). There was >97% overlap of native tdT expression with immunoreactivity to tdT using the DSRed antibody. tdT^+^ neurons were significantly smaller than tdT^−^ neurons in both ganglia (p<0.05, Fig. 2C). We then sought to correlate tdT expression in sensory neurons from *TRPV1^Flp/+^R26^ai65f/+^* mice with sensitivity to the selective TRPV1 agonist capsaicin. Similar to the intact ganglia, 38.8% (33/85) of dissociated vagal neurons and 29.2% (42/144) of dissociated DRG neurons expressed tdT (Fig. 2D). Capsaicin (1μM) evoked a robust increase in [Ca^2+^]_cyt_ in vagal tdT^+^ neurons (mean of 0.53 ± 0.05) compared to vagal tdT^−^ neurons (mean of 0.18 ± 0.04, p<0.05), and in DRG tdT^+^ neurons (mean of 0.82 ± 0.06) compared to DRG tdT^−^ neurons (mean of 0.16 ± 0.03, p<0.05) (Fig. 2E). Mean responses to the P2X_2/3_ agonist αβmATP (10μM) were also significantly greater in vagal tdT^+^ neurons compared to vagal tdT^−^ neurons (0.16 ± 0.02 vs. 0.11 ± 0.02, p<0.05) and in DRG tdT^+^ neurons compared to DRG tdT^−^ neurons (0.02 ± 0.004 vs. 0.01 ± 0.001, p<0.05), but such differences were limited compared to the difference in αβmATP responses between vagal and DRG neurons (p<0.05). Analysis of individual responses showed that 100% (33/33) of vagal tdT^+^ neurons and 92.9% (39/42) of DRG tdT^+^ neurons were activated by capsaicin compared to only 40.4% (21/52) of tdT^−^ vagal neurons and 30.4% (31/102) of tdT^−^ DRG neurons (Fig. 2F). As such, 33 out of the 54 (61.1%) vagal capsaicin-sensitive neurons were tdT^+^, and 39 out of the 70 (55.7%) DRG capsaicin-sensitive neurons were tdT^+^. Many vagal neurons (70.1%, 60/85) were also activated by αβmATP, compared to 6.9% (10/144) of DRG neurons (Fig. 2F). Overall, the data from *TRPV1^Flp/+^R26^ai65f/+^*mice suggests that this reporter selectively labels TRPV1-expressing sensory neurons, although not with full efficiency.

**Figure 2:**
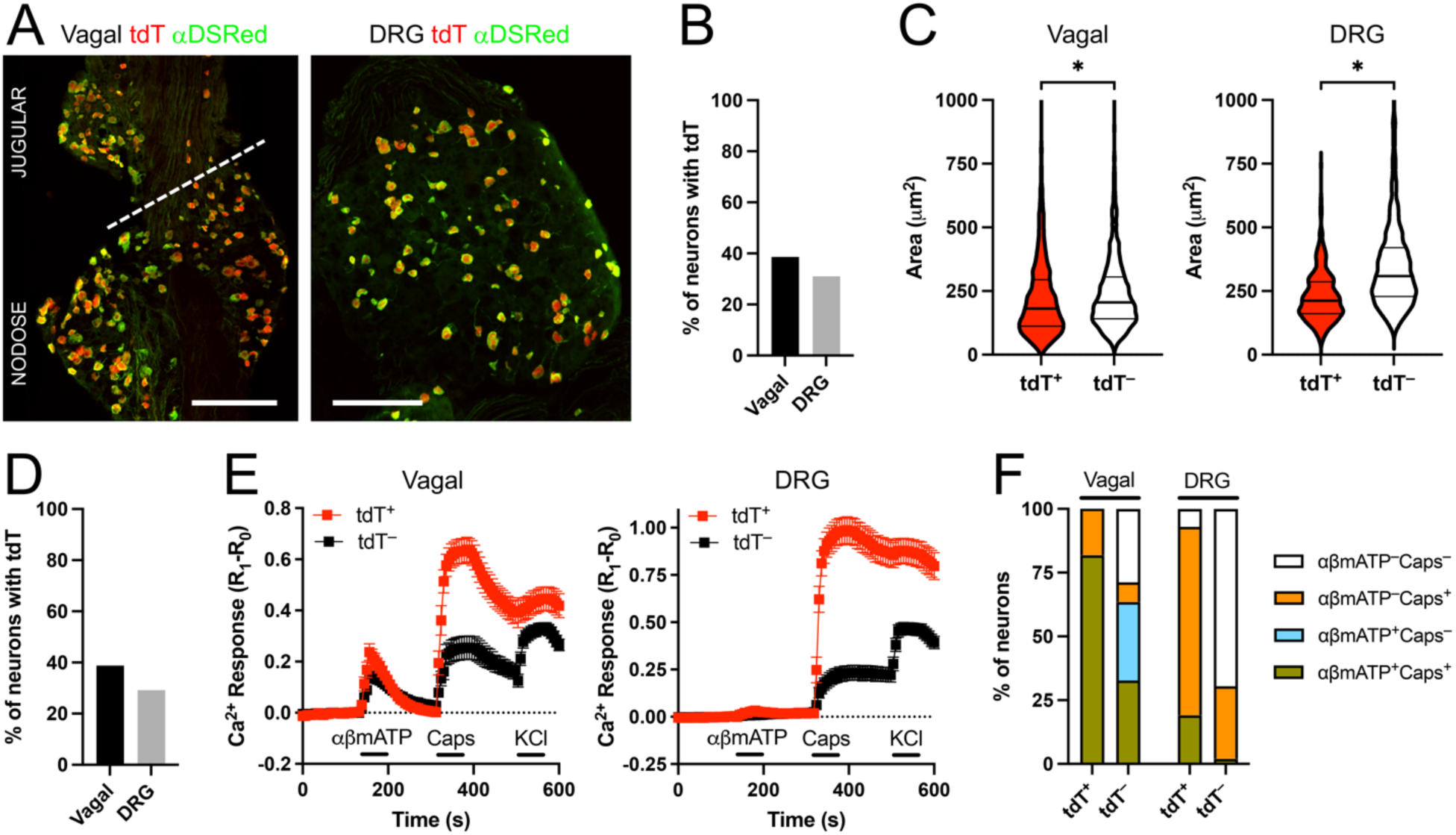
Subsets of capsaicin-sensitive sensory neurons in *TRPV1^Flp^R26^ai65f^*mice express tdTomato. A, representative frozen sections of native tdTomato (tdT) expression (in red) compared with immunoreactivity to tdT (DsRed, in green) in vagal ganglia (left) and thoracic DRG (right). Scale bars denotes 200μm. B, percentage of neurons in frozen sections of the vagal ganglia and DRG (n=4243 and 1502 neurons, respectively) that express tdT. C, violin plots of the cell body area (in μm^2^) of tdT^+^ and tdT^−^ neurons in the vagal ganglia (left) and thoracic DRG (right). * denotes significant difference in cell body area (p<0.05). D, percentage of dissociated vagal and thoracic DRG neurons (n=85 and 144, respectively) that express tdT. E, mean ± SEM 340/380 ratio [Ca^2+^]_i_ responses of dissociated vagal neurons (left, n of 33 tdT^+^ (in red), and n of 52 tdT^−^ (in black)) and thoracic DRG neurons (right, n of 42 tdT^+^, and n of 102 tdT^−^) to αβmATP (10μM), capsaicin (Caps, 1μM) and KCl (75mM). F, percentage of tdT^+^ and tdT^−^ neurons dissociated from the vagal ganglia and thoracic DRG that were activated by AITC and capsaicin.

We further investigated the Flp efficiency in vagal neurons using a series of crosses of *TRPV1^Flp^* mice with the Flp-sensitive reporter strain *R26^FLTG^*(which expresses tdTomato in Flp-expressing cells) to produce mice that were either homozygous or heterozygous for both alleles. Frozen sections of vagal ganglia showed progressive increases in the percentage of neurons expressing tdT from *TRPV1^Flp/+^R26^FLTG/+^* mice to *TRPV1^Flp/Flp^R26^FLTG/FLTG^* mice (Fig. 3), with homozygosity for the Flp allele appearing to play a determinant role. This suggests that Flp expression is rate-limiting for the recombination of the ROSA26 reporters in these sensory neurons. Additionally, the efficiency of vagal sensory neuron tdT expression in *TRPV1^Flp/+^R26^FLTG/+^*mice (Fig. 3B) was reduced compared to *TRPV1^Flp/+^R26^ai65f/+^* mice (Fig. 2B), suggesting that the larger *R26^FLTG^* insert was less efficiently recombined.

**Figure 3:**
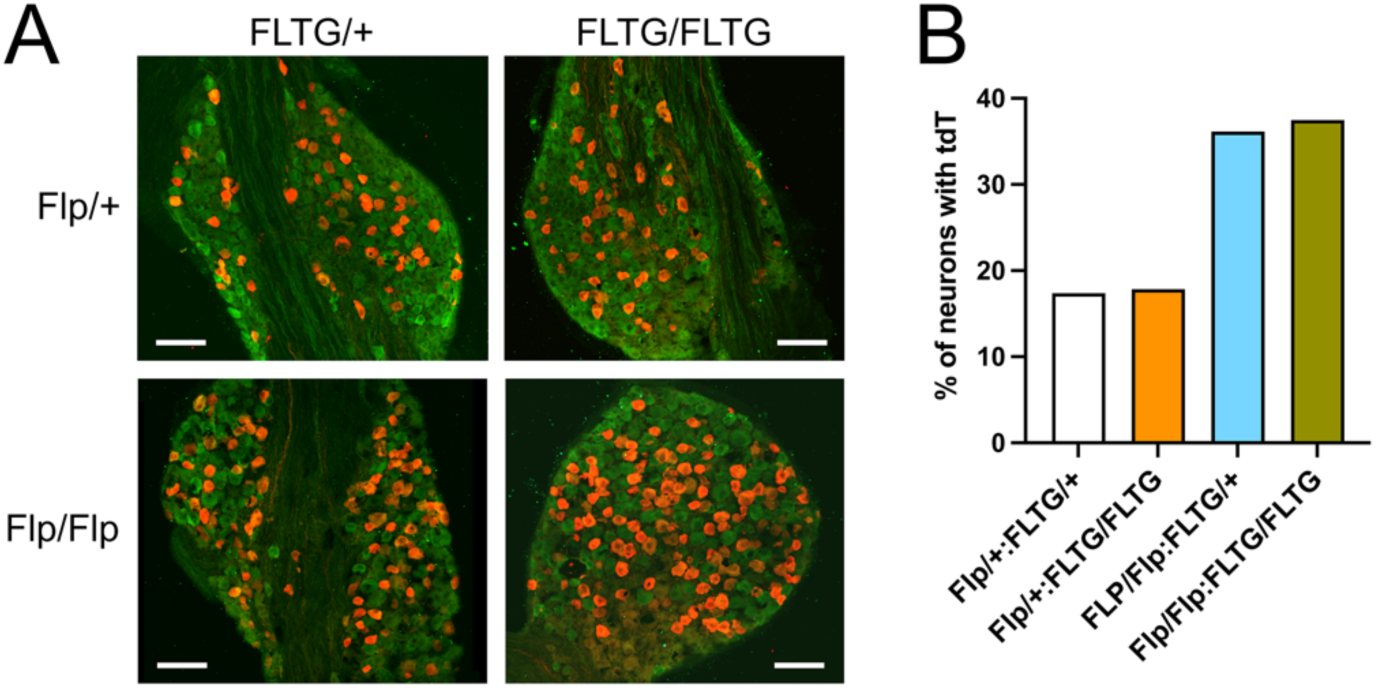
Efficiency of tdTomato expression in vagal sensory neurons in *TRPV1^Flp^R26^FLTG^*mice. Representative frozen sections (A) and quantification (B) of tdT expression (in red) and the pan neuronal marker pgp9.5 (in green) in vagal ganglia from *TRPV1^Flp/+^R26^FLTG/+^* mice (3206 neurons), *TRPV1^Flp/+^R26^FLTG/FLTG^* mice (2634 neurons), *TRPV1^Flp/Flp^R26^FLTG/+^* mice (5744 neurons) and *TRPV1^Flp/Flp^R26^FLTG/FLTG^*mice (7396 neurons). Scale bars denotes 100μm.

### Intersectional labeling of nociceptive subsets within the vagal ganglia

We chose to use the *R26^FLTG^*strain, despite its limited efficiency, for an intersectional approach to label genetically-distinct nociceptive subsets, because its reporter is sensitive to both Cre and Flp: producing GFP in Flp^+^Cre^+^ cells and tdT in Flp^+^Cre^−^ cells.

The purinergic ion channel P2X_2_ is selectively expressed in nodose neurons but not jugular neurons ^3,27,35,38,39^. Through a series of crosses of *TRPV1^Flp^* mice, *P2X2^Cre^* mice and *R26^FLTG^* mice we generated heterozygous *TRPV1^Flp^P2X2^Cre^-FLTG* mice that were expected to express GFP in TRPV1^+^P2X2^+^ neurons and tdT in TRPV1^+^P2X2^−^ neurons. Frozen sections of the vagal ganglia (n=5) from these mice showed GFP and tdT expression in distinct neurons of which 75.0% (120/160) and 82.8% (101/122), respectively, also displayed immunoreactivity for TRPV1 (Fig. 4A, F). Although the nodose and jugular vagal ganglia are partially fused in mice, it is nonetheless possible to approximate the division of the ganglia anatomically ^7,27^. We found that GFP expression in *TRPV1^Flp^P2X2^Cre^-FLTG* mice was restricted to a subset of nodose neurons (15.0%, 160/1066) and was absent in jugular neurons (0%, 0/1367), whereas tdT expression was detected in 4.4% (47/1066) and 5.5% (75/1367) of nodose and jugular neurons, respectively (Fig. 4).

**Figure 4:**
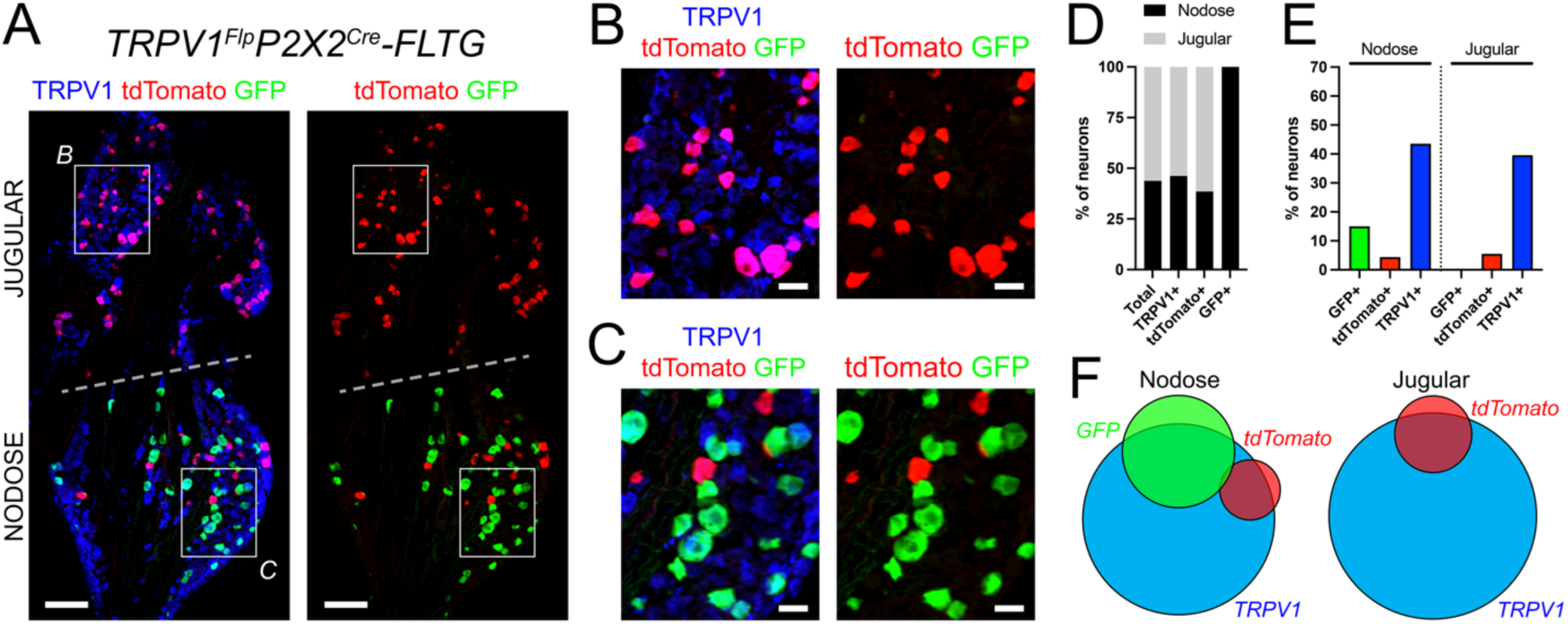
Intersectional labeling of distinct nociceptive vagal subsets using *TRPV1^Flp^P2X2^Cre^-FLTG* mice. A, representative vagal ganglion frozen section showing expression of tdTomato (in red) and GFP (in green) with (left) and without (right) immunoreactivity to TRPV1 (in blue). B, higher magnification of jugular area identified by white box in A. C, higher magnification of nodose area identified by white box in A. D, comparative percentage contribution of nodose and jugular neurons to the total vagal neuron population (n=2433), and the TRPV1^+^ (n=1005), GFP^+^ (n=160) and tdT^+^ (n=122) subsets. E, percentage of nodose (n=1066) and jugular (n=1367) neurons that express GFP, tdTomato or TRPV1. F, Euler diagram showing the overlap of nodose neurons (left) and jugular neurons (right) expressing GFP, tdTomato and TRPV1. Scale bars denote 100μm (A), 25μm (B, C).

Tac1, the gene for that encodes substance P and other tachykinins, is expressed in jugular TRPV1^+^ neurons as well as some nodose TRPV1^−^ neurons ^3,7^. Through a series of crosses of *TRPV1^Flp^* mice, *Tac1^Cre^* mice and *R26^FLTG^* mice we generated heterozygous *TRPV1^Flp^Tac1^Cre^-FLTG* mice that were expected to express GFP in TRPV1^+^Tac1^+^ neurons and tdT in TRPV1^+^Tac1^−^ neurons. Frozen sections of the vagal ganglia (n=10) from these mice showed GFP and tdT expression in distinct neurons of which 86.2% (313/363) and 85.3% (406/476), respectively, also expressed TRPV1 (Fig. 5A, F). We found GFP was preferentially expressed in jugular neurons (11.8%, 301/2560) compared to nodose neurons (1.6%, 62/3797) (Fig. 5). Whereas tdT was preferentially expressed in a subset of nodose neurons (12.2%, 463/3797) compared to jugular neurons (0.5%, 13/2560)(Fig. 5).

**Figure 5:**
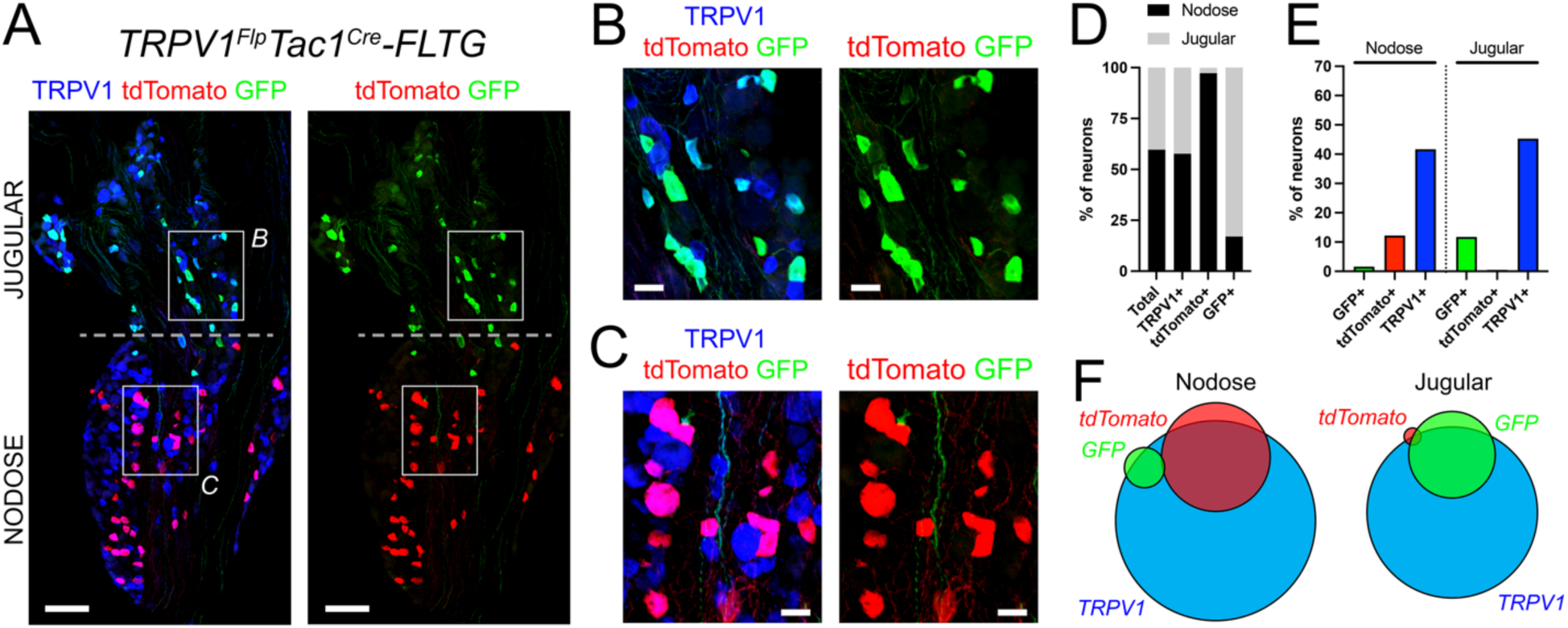
Intersectional labeling of distinct nociceptive vagal subsets using *TRPV1^Flp^Tac1^Cre^-FLTG* mice. A, representative vagal ganglion frozen section showing expression of tdTomato (in red) and GFP (in green) with (left) and without (right) immunoreactivity to TRPV1 (in blue). B, higher magnification of jugular area identified by white box in A. C, higher magnification of nodose area identified by white box in A. D, comparative percentage contribution of nodose and jugular neurons to the total vagal neuron population (n=6357), and the TRPV1^+^ (n=2741), GFP^+^ (n=363) and tdT^+^ (n=476) subsets. E, percentage of nodose (n=3797) and jugular (n=2560) neurons that express GFP, tdTomato or TRPV1. F, Euler diagram showing the overlap of nodose neurons (left) and jugular neurons (right) expressing GFP, tdTomato and TRPV1. Scale bars denote 100μm (A), 25μm (B, C).

We next sought to correlate tdT and GFP expression with evidence of TRPV1, P2X_2_ and Tac1 expression. We first performed single cell RT-PCR on 50 dissociated vagal *TRPV1^Flp^P2X2^Cre^-FLTG* neurons (from 6 heterozygous mice) and 51 dissociated vagal *TRPV1^Flp^Tac1^Cre^-FLTG* neurons (from 6 heterozygous mice), all but two of which expressed the vesicular glutamate transporter 2 (VGLUT2) (Fig. 6A, B). We found TRPV1 transcript in 100% of GFP^+^ (26/26) and tdT^+^ (24/24) neurons from *TRPV1^Flp^P2X2^Cre^-FLTG* vagal ganglia, consistent with the requirement of TRPV1 expression for reporter expression. P2X_2_ transcript expression was found in 92.3% (24/26) of GFP^+^ neurons and in 50.0% (12/24) of tdT^+^ neurons, whereas Tac1 expression was found in 7.7% (2/26) of GFP^+^ neurons and in 70.8% (17/24) of tdT^+^ neurons. TRPV1 transcript was found in 96% of GFP^+^ (25/26) and tdT^+^ (24/25) neurons from *TRPV1^Flp^Tac1^Cre^-FLTG* vagal ganglia. Tac1 transcript expression was found in 100% (26/26) of GFP^+^ neurons and in 12.0% (3/25) of tdT^+^ neurons, whereas P2X_2_ expression was found in 19.2% (5/26) of GFP^+^ neurons and in 96.0% (24/25) of tdT^+^ neurons. There was limited overlap of P2X_2_ and Tac1 expression in the GFP^+^ or tdT^+^ neurons from either strain of mice (Fig. 6C).

**Figure 6:**
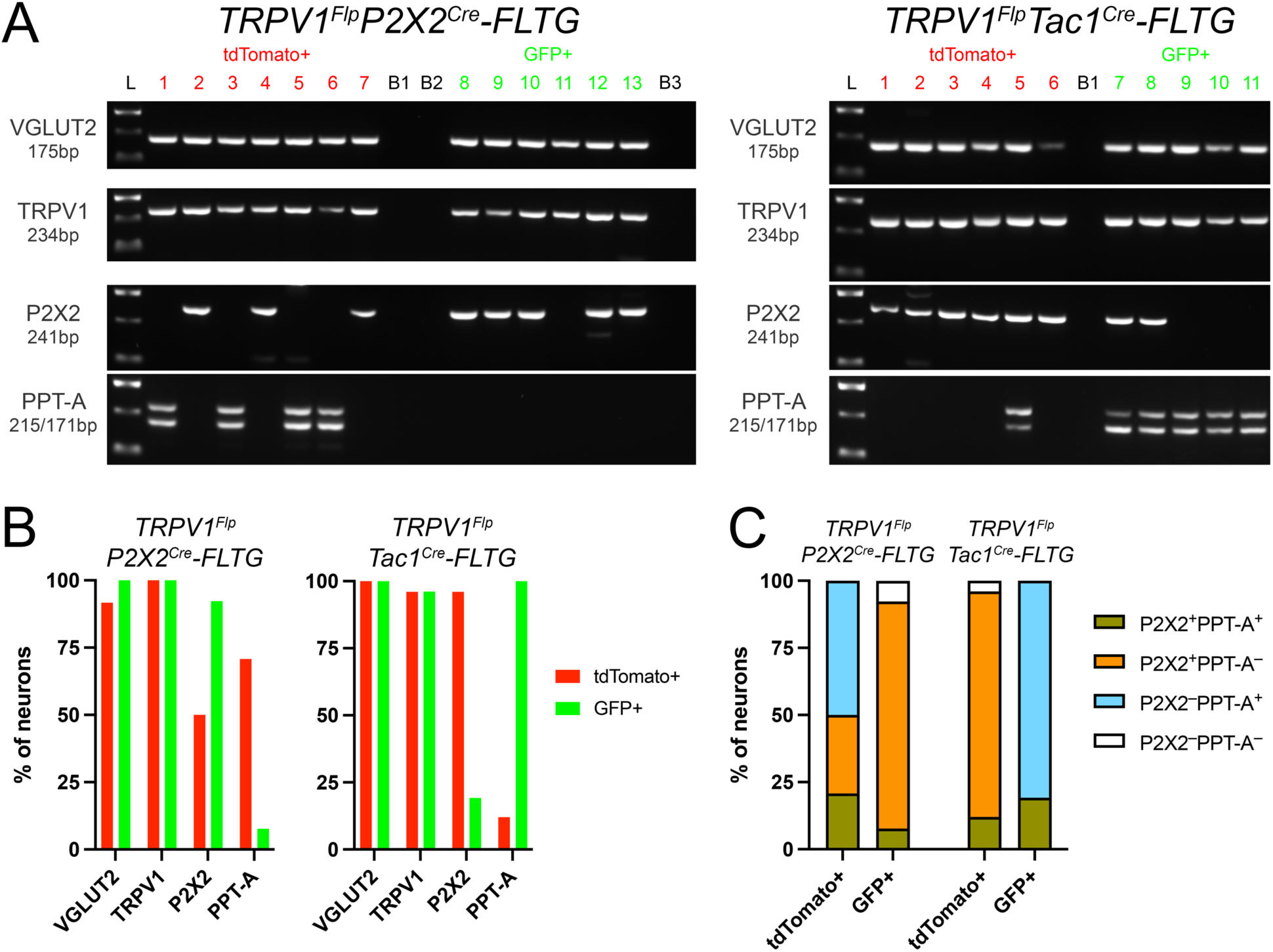
Intersectional reporter expression correlates with the transcript signature of distinct vagal afferent subsets. A, representative gels of individual neurons dissociated from the vagal ganglia of *TRPV1^Flp^P2X2^Cre^-FLTG* mice (left; 1-7 tdT^+^, 8-13 GFP^+^) and *TRPV1^Flp^Tac1^Cre^-FLTG* mice (right; 1-6 tdT^+^, 7-11 GFP^+^) following RT-PCR for transcripts for VGLUT2 (175 bp), TRPV1 (234 bp), P2X2 (241 bp), and PPT-A (215/171/162/118 bp). Also shown Hyperladder 100bp (L) and negative bath controls (B1 to B3). B, percentage of tdT^+^ (in red) and GFP^+^ (in green) neurons from *TRPV1^Flp^P2X2^Cre^-FLTG* vagal ganglia (left, n=24 and 26, respectively) and *TRPV1^Flp^Tac1^Cre^-FLTG* vagal ganglia (right, n=25 and 26, respectively) that express VGLUT2, TRPV1, P2X2, and PPT-A. C, overlap of P2X2 and PPT-A expression in tdT^+^ and GFP^+^ vagal neurons from *TRPV1^Flp^P2X2^Cre^-FLTG* mice (left) and *TRPV1^Flp^Tac1^Cre^-FLTG* mice (right).

We then investigated neuronal sensitivity to αβmATP (10μM) and capsaicin (1μM) using Fura2AM Ca^2+^ imaging. Expression of GFP and tdT was found in 14.5% (94/649) and 10.0% (65/649) of dissociated vagal neurons, respectively, from heterozygous *TRPV1^Flp^P2X2^Cre^-FLTG* mice (3 animals)(Fig. 7A, B). Capsaicin evoked greater increases in [Ca^2+^]_cyt_ in GFP^+^ neurons (mean of 0.57 ± 0.04, p<0.05) and in tdT^+^ neurons (mean of 0.86 ± 0.05, p<0.05) compared to reporter-negative neurons (mean of 0.20 ± 0.02) (Fig. 7C). Whereas αβmATP evoked substantial increases in [Ca^2+^]_cyt_ in GFP^+^ neurons (mean of 0.27 ± 0.02) compared to tdT^+^ neurons (mean of 0.05 ± 0.02, p<0.05) (Fig. 7C). αβmATP-evoked responses in reporter-negative neurons (mean of 0.15 ± 0.01) were significantly greater than those of tdT^+^ neurons (p<0.05), but less than those of GFP^+^ neurons (p<0.05). Analysis of individual responses showed that 90.4% (85/94) of GFP^+^ neurons were αβmATP^+^capsaicin^+^ with the remainder being αβmATP^−^capsaicin^+^, whereas only 23.1% (15/65) of tdT^+^ neurons were αβmATP^+^capsaicin^+^ with the remainder being αβmATP^−^ Capsaicin^+^ (Fig. 7D).

**Figure 7:**
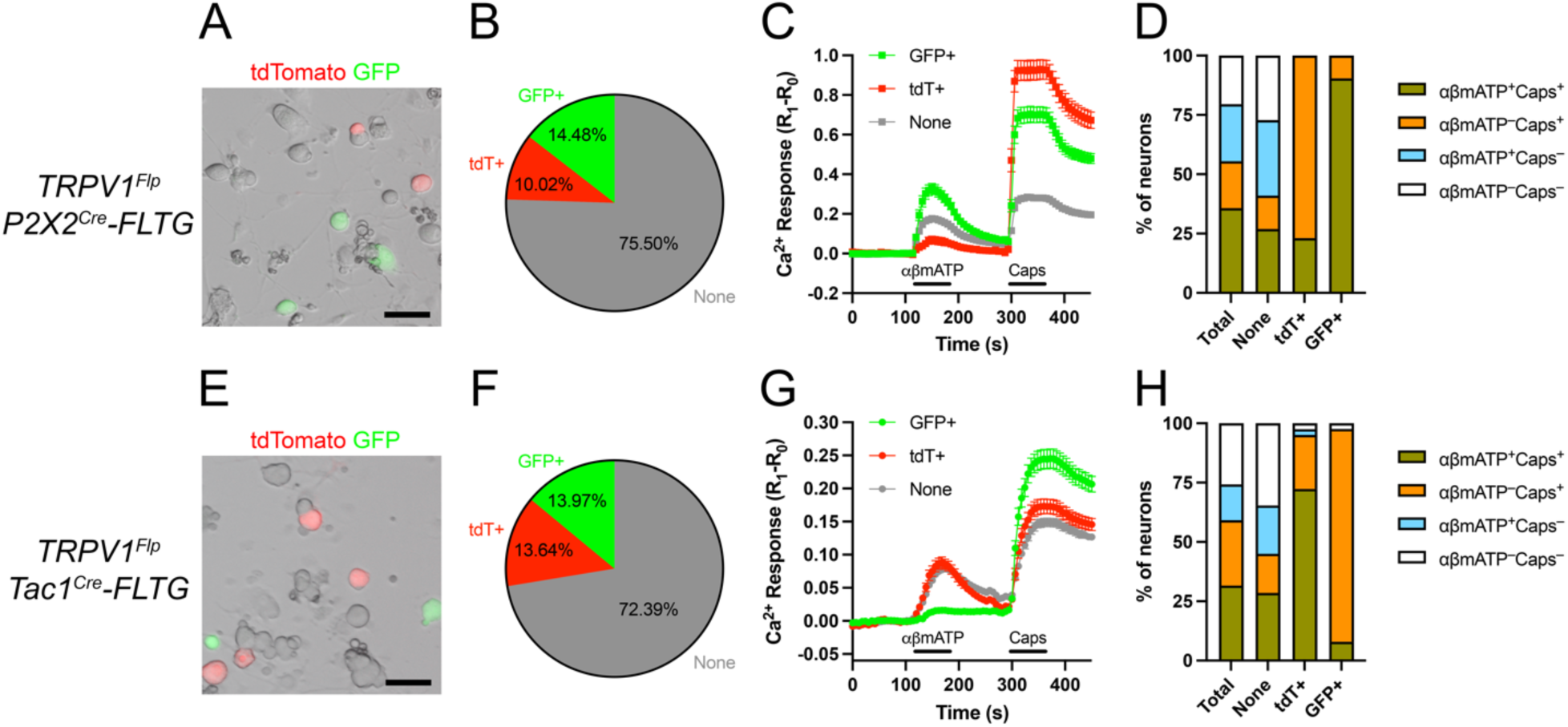
Intersectional reporter expression correlates with the functional signature of distinct vagal afferent subsets. A-D, *TRPV1^Flp^P2X2^Cre^-FLTG* vagal neurons (n=649). E-H, *TRPV1^Flp^Tac1^Cre^-FLTG* vagal neurons (n=902). A and E, representative brightfield image of dissociated neurons with overlay of tdTomato (in red) and GFP (in green) expression. Scale bars denotes 50μm. B and F, percentage of neurons expressing tdTomato and GFP. C and G, mean ± SEM 340/380 ratio [Ca^2+^]_i_ responses of tdT^+^, GFP^+^ or reporter-negative (None) vagal neurons to αβmATP (10μM) and capsaicin (Caps, 1μM). D and H, percentage of the total vagal neuron population, and the reporter-negative (None), GFP^+^ and tdT^+^ subsets that were activated by AITC and capsaicin.

Following the dissociation of vagal ganglia from heterozygous *TRPV1^Flp^Tac1^Cre^-FLTG* mice (3 animals), GFP and tdT expression was detected in 14.0% (126/902) and 13.6% (123/902) of neurons, respectively (Fig. 7E, F). Capsaicin evoked greater increases in [Ca^2+^]_cyt_ in GFP^+^neurons (mean of 0.18 ± 0.01, p<0.05) and in tdT^+^ neurons (mean of 0.11 ± 0.01, p<0.05) compared to reporter-negative neurons (mean of 0.08 ± 0.005) (Fig. 7G). Whereas αβmATP evoked substantial increases in [Ca^2+^]_cyt_ in tdT^+^ neurons (mean of 0.06 ± 0.01, p<0.05) and reporter-negative neurons (mean of 0.06 ± 0.003), compared to the minimal responses in GFP^+^ neurons (mean of 0.01 ± 0.003) (Fig. 7G). Analysis of individual responses showed that only 7.9% (10/126) of GFP^+^ neurons were αβmATP^+^capsaicin^+^ with 89.7% (113/126) being αβmATP^−^ capsaicin^+^, whereas 72.4% (89/123) of tdT^+^ neurons were αβmATP^+^capsaicin^+^ with 22.8% (28/123) being αβmATP^−^capsaicin^+^ (Fig. 7H).

The lack of TRPV1 immunoreactivity in 17-35% of reporter^+^ neurons in the vagal ganglia of *TRPV1^Flp^Tac1^Cre^-FLTG* and *TRPV1^Flp^Tac1^Cre^-FLTG* mice (Fig. 4F, 5F) could suggest reporter expression was triggered by transient TRPV1 expression earlier in development. However, it is likely that TRPV1 immunoreactivity has been underreported here (due to antibody inefficiencies) given that TRPV1 transcript (Fig. 6B) and capsaicin-sensitivity (Fig. 7D, H) was observed in 98.0% and 97.8% of reporter^+^ neurons, respectively.

We also mapped vagal nociceptive neurons using dual recombinase-sensitive mCherry-expressing AAV, microinjected into the vagal ganglia. This intersectional approach drives mCherry expression dependent on the active expression of Cre and Flp, and thus is not affected by any transient recombinase expression earlier in development. Intravagal injection of homozygous *TRPV1^Flp^P2X2^Cre^* mice with AAV9-ConFon-mCherry evoked mCherry expression in 510 out of 2255 vagal neurons (n of 3 animals), 97% of which were located in the nodose ganglia (Fig. 8A, E, F), consistent with GFP expression in vagal TRPV1^+^P2X2^+^ neurons in *TRPV1^Flp^P2X2^Cre^-FLTG* mice. Intravagal AAV9-ConFon-mCherry injection in homozygous *TRPV1^Flp^Tac1^Cre^*mice evoked mCherry expression in 429 out of 5066 vagal neurons (n of 4 animals), 96% of which were found in the jugular ganglia (Fig. 8B, E, F), consistent with GFP expression in vagal TRPV1^+^Tac1^+^ neurons in *TRPV1^Flp^Tac1^Cre^-FLTG* mice. Intravagal AAV9-CoffFon-mCherry injection in homozygous *TRPV1^Flp^Tac1^Cre^* mice evoked mCherry expression in 390 out of 1875 vagal neurons (n of 3 animals), 85% of which were found in the nodose ganglia (Fig. 8C, E, F). Tac1^−^TRPV1^+^ neurons are likely to include nodose P2X2^+^ neurons and jugular P2X2^−^ neurons. Lastly, intravagal AAV9-ConFoff-mCherry injection in homozygous *TRPV1^Flp^Tac1^Cre^*mice evoked mCherry expression in 575 out of 2606 vagal neurons (n of 3 animals), 59% of which were found in the jugular ganglia (Fig. 8D, E, F).

**Figure 8:**
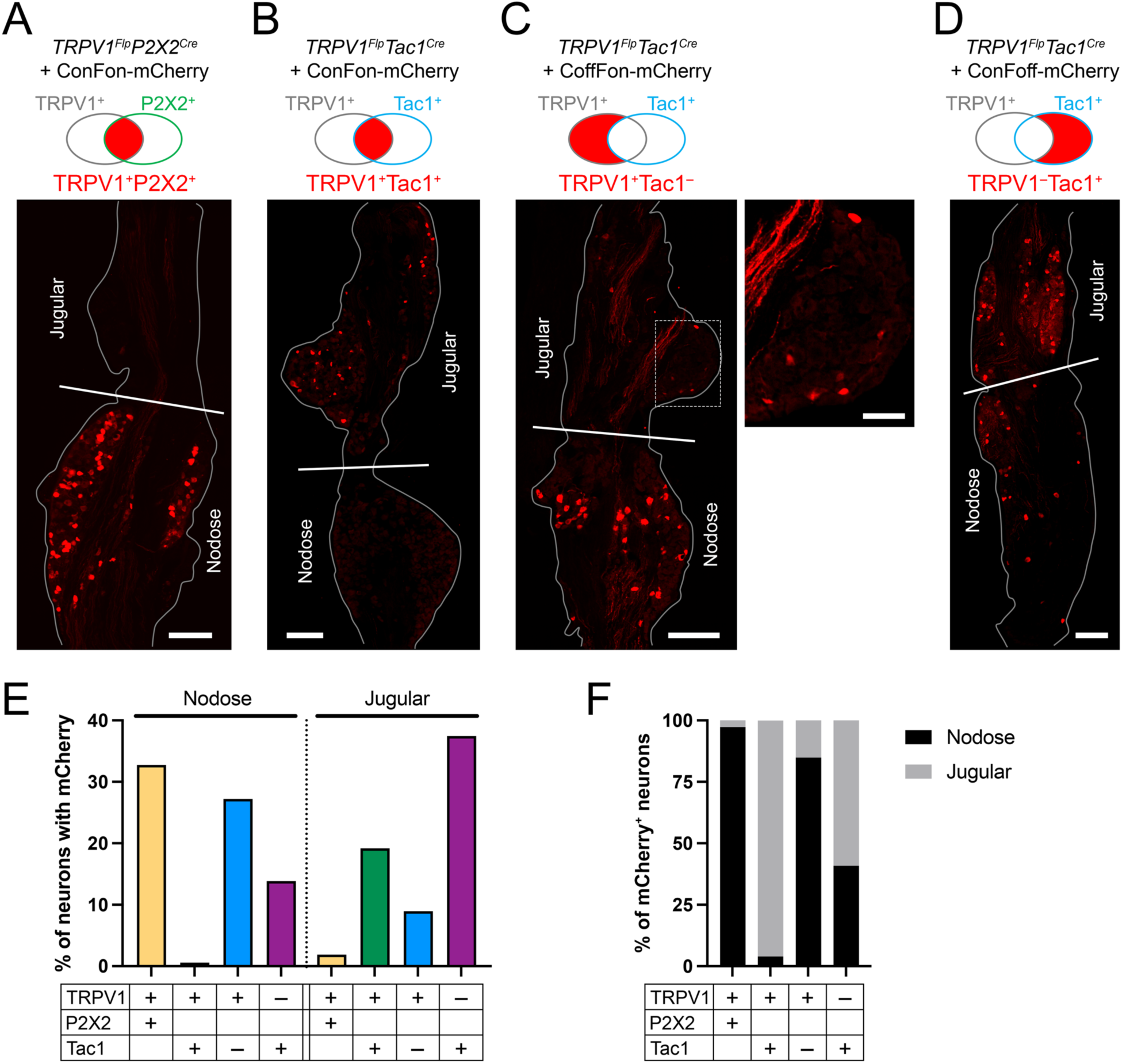
Intersectional labeling of distinct vagal subsets following intravagal injection with AAV9. A-D, representative vagal ganglion frozen sections showing expression of mCherry (in red) following injection of AAV9-ConFon-mCherry into *TRPV1^Flp^P2X2^Cre^* mice (A) and *TRPV1^Flp^Tac1^Cre^* mice (B), AAV9-CoffFon-mCherry into *TRPV1^Flp^Tac1^Cre^* mice (C), and AAV9-ConFoff-mCherry into *TRPV1^Flp^Tac1^Cre^* mice (D). Grey line denotes ganglion outline. White line indicates the approximate division of nodose and jugular portions of the vagal ganglion. White dotted square identifies a region shown in higher magnification in insert to the right of C. Scale bars denote 150μm, and 50μm in insert. E, percentage of nodose and jugular neurons that express mCherry following AAV9 injection. F, the percentage distribution of mCherry+ neurons in the nodose and jugular ganglion.

### Distinct central terminations of vagal nociceptive subsets

Previous studies have identified the nTS, area postrema (AP) and Pa5 as the targets of central terminations of vagal afferents ^7,9,27,28,40^. We found dense GFP^+^ fibers within the dorsal and medial subnuclei of the caudal nTS of heterozygous *TRPV1^Flp^P2X2^Cre^-FLTG* mice (n=3 animals, Fig. 9A). GFP^+^ fibers also innervated the AP, but not the Pa5. Whereas tdT^+^ fibers densely innervated the Pa5, with very sparse labeling in the nTS subnuclei and none in the AP (Fig. 9A). In the medulla of heterozygous *TRPV1^Flp^Tac1^Cre^-FLTG* mice (n=3 animals), GFP^+^ fibers densely innervated the Pa5, with almost no innervation of the nTS or AP (Fig. 9B). Whereas tdT^+^ fibers innervated the dorsal and medial subnuclei of the caudal nTS, the AP, and the Pa5 (Fig. 9B). Combined these data suggest that nodose TRPV1^+^P2X2^+^ afferents innervate the nTS (and to a lesser extent AP), and jugular TRPV1^+^Tac1^+^ afferents innervate the Pa5. The tdT^+^ fibers in the Pa5 of *TRPV1^Flp^Tac1^Cre^-FLTG* mice (Fig. 9B) suggest that some TRPV1^+^Tac1^−^ afferents also innervate this region but, given the lack of GFP^+^ fibers within the Pa5 of *TRPV1^Flp^P2X2^Cre^-FLTG* mice (Fig. 9A), these fibers do not express P2X2.

**Figure 9:**
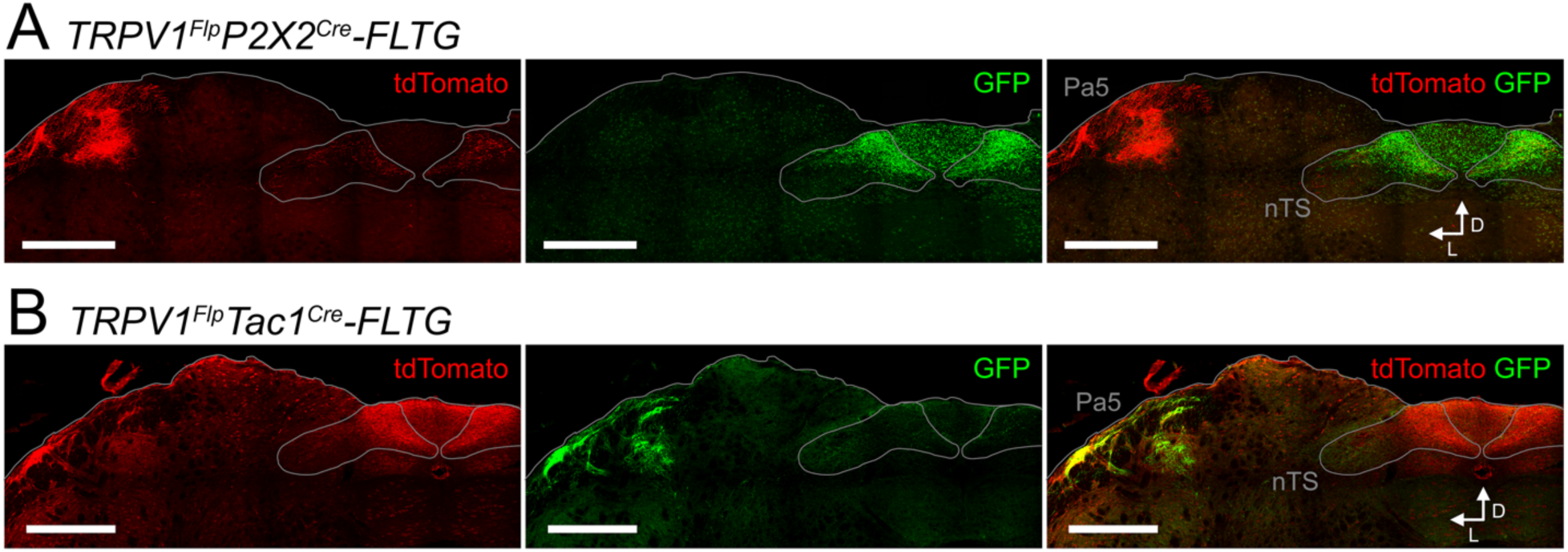
Distinct brainstem terminations of nociceptive afferent subsets. A and B, tdTomato^+^ and GFP^+^ fibers innervating the dorsal medulla (coronal slice 300μm caudal to obex) in the *TRPV1^Flp^P2X2^Cre^-FLTG* (A) and *TRPV1^Flp^Tac1^Cre^-FLTG* (B) mouse. Scale bars denote 500μm. Grey lines denote the medulla outline and the nTS. White arrows denote dorsal aspect and lateral aspect from midline.

Given that the TRPV1^Flp^ allele also drives FLTG-dependent reporter expression in trigeminal afferents (not shown), it is possible that some of the reporter^+^ afferents in the medulla of *TRPV1^Flp^P2X2^Cre^-FLTG* or *TRPV1^Flp^Tac1^Cre^-FLTG* mice are non-vagal. To specifically map central terminations of vagal afferents, we performed unilateral vagal microinjection with intersectional mCherry-expressing AAV9. Following injection of homozygous *TRPV1^Flp^P2X2^Cre^* mice with AAV9-ConFon-mCherry (n=3), we found dense ipsilateral and partial contralateral innervation of the dorsal and medial subnuclei of the caudal nTS by mCherry^+^ fibers (Fig. 10A). mCherry^+^ fibers also innervated the AP, but innervation of the Pa5 was almost absent. Whereas we found dense mCherry^+^ innervation of the Pa5 (only ipsilateral) but not the nTS or AP following AAV9-ConFon-mCherry injection of homozygous *TRPV1^Flp^Tac1^Cre^* mice (n=3 animals, Fig. 10B). These data confirm that vagal TRPV1^+^P2X2^+^ afferents exclusively innervate the nTS and AP, and vagal TRPV1^+^Tac1^+^ afferents exclusively innervate the Pa5. Following injection of homozygous *TRPV1^Flp^Tac1^Cre^*mice with AAV9-CoffFon-mCherry (n=3), we found dense ipsilateral and partial contralateral TRPV1^+^Tac1^−^ innervation of the dorsal and medial subnuclei of the caudal nTS and AP (Fig. 10C). We also found evidence of vagal TRPV1^+^Tac1^−^ innervation of the ipsilateral Pa5. Lastly, we found vagal TRPV1^−^Tac1^+^ fibers innervated the ipsilateral Pa5 and the lateral subnuclei of the nTS (mainly ipsilateral) following AAV9-ConFoff-mCherry injection of homozygous *TRPV1^Flp^Tac1^Cre^*mice (n=3 animals, Fig. 10D).

**Figure 10:**
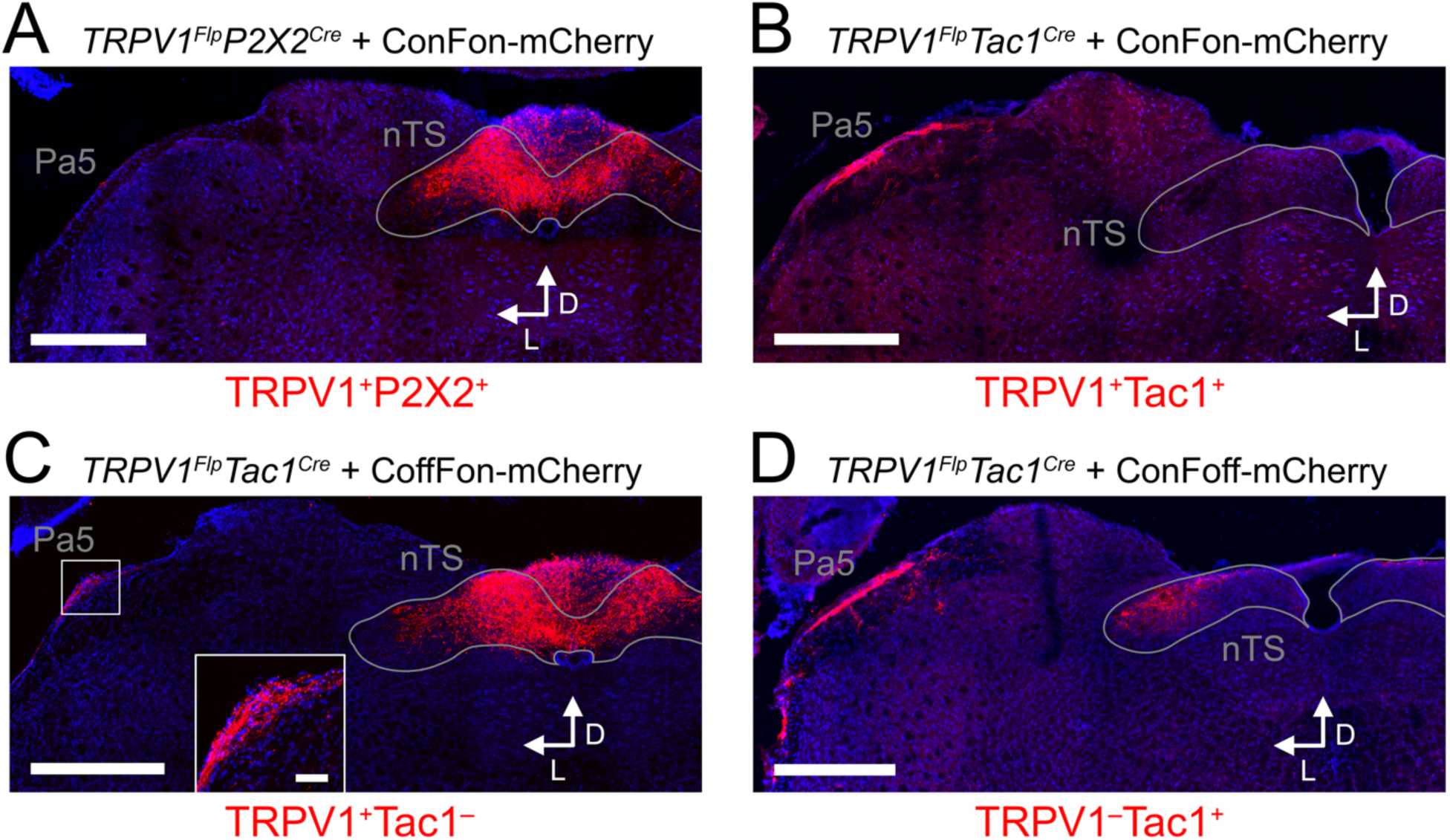
Distinct brainstem terminations of vagal afferent subsets. Representative coronal sections of the dorsal medulla showing expression of mCherry (in red) following unilateral vagal injection of AAV9-ConFon-mCherry into *TRPV1^Flp^P2X2^Cre^* mice (A, 240μm caudal to obex) and *TRPV1^Flp^Tac1^Cre^*mice (B, 30μm caudal to obex), AAV9-CoffFon-mCherry into *TRPV1^Flp^Tac1^Cre^* mice (C, 240μm caudal to obex), and AAV9-ConFoff-mCherry into *TRPV1^Flp^Tac1^Cre^* mice (D, 30μm caudal to obex). DAPI staining (in blue) also shown. Grey lines denote the outline of the nTS. White arrows denote dorsal aspect and lateral aspect from midline. In C, a high magnification of the Pa5 is shown in the insert. Scale bars denote 500μm, and 50μm in insert.

### Distinct innervation of the lungs by vagal nociceptive subsets

Vagal innervation supplies the majority of sensory fibers to the lower airways ^10^. Previous studies have shown vagal TRPV1^+^ fibers innervate most conducting airways throughout the bronchial tree as well as a small proportion of pulmonary blood vessels ^9,10^. TRPV1^+^ fibers also project from conducting airways into the alveolar region. Here, we used intersectional approaches to map the innervation of the lungs by TRPV1^+^P2X2^+^ and TRPV1^+^Tac1^+^ afferents.

In the lungs of heterozygous *TRPV1^Flp^P2X2^Cre^-FLTG* mice (n=5 animals), we found GFP^+^ fibers innervating 65% of conducting airways and 10% of pulmonary blood vessels (Fig. 11A, C, E). GFP^+^ fibers also projected from 36% of conducting airways into the alveolar region (Fig. 11A, D). Whereas tdT^+^ fibers innervated 80% of conducting airways but only projected into the alveolar region from 8% of conducting airways (Fig. 11A, C, D). tdT^+^ fibers also innervated 18% of blood vessels (Fig. 11A, E). In the lungs of heterozygous *TRPV1^Flp^Tac1^Cre^-FLTG* mice (n=6 animals), we found GFP^+^ fibers innervating 80% of conducting airways and 15% of pulmonary blood vessels (Fig. 11B, C, E). Importantly, no GFP^+^ fibers projected from the conducting airways into the alveolar region (Fig. 11B, D). Whereas 61% of conducting airways were innervated by tdT^+^ fibers, almost half of which (27%) were innervated by tdT^+^ fibers that projected into the alveolar region (Fig. 11B-D). tdT^+^ fibers also innervated 9% of blood vessels (Fig. 11B, E).

**Figure 11:**
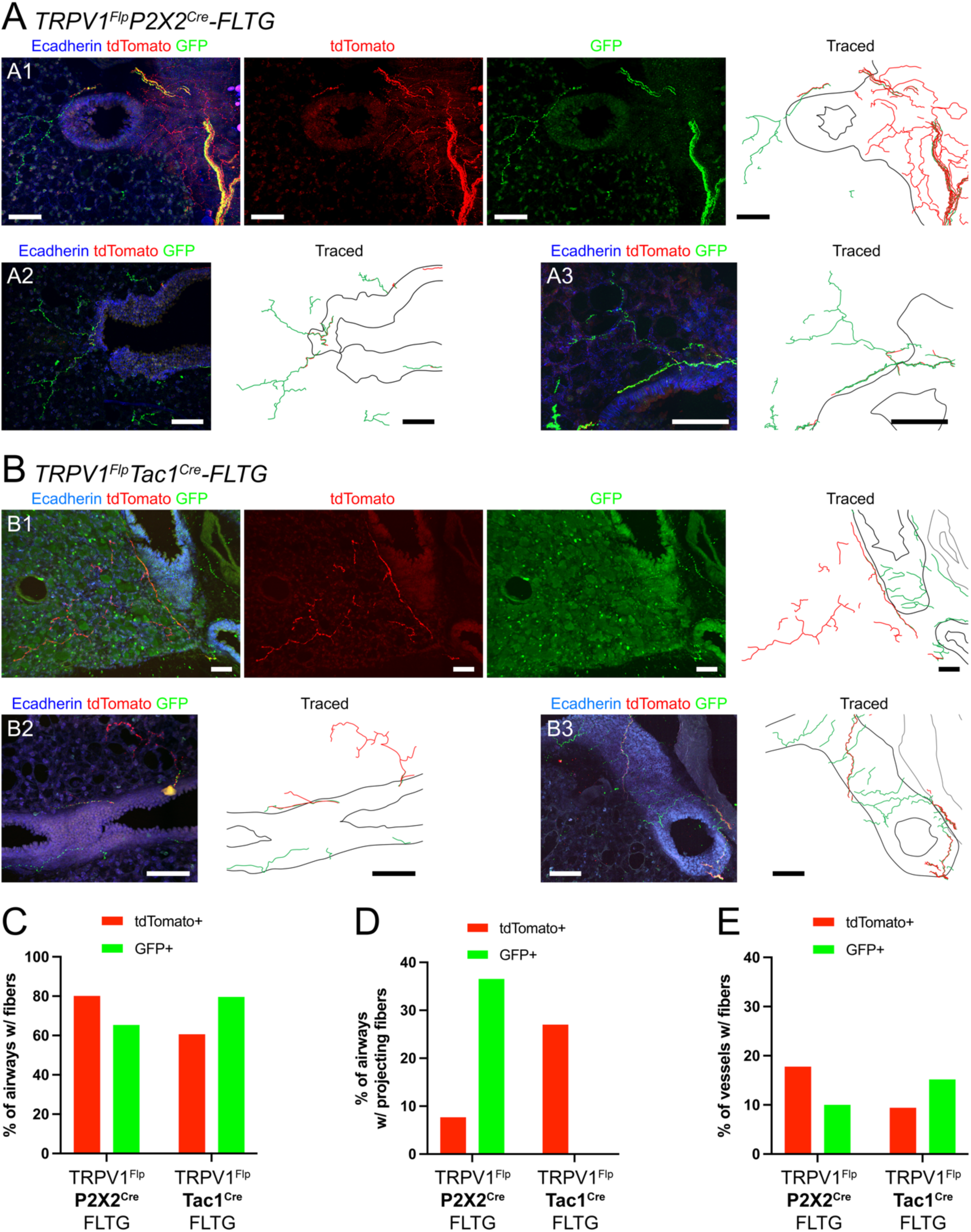
Innervation of the lung by nociceptive afferent subsets. A and B, tdTomato^+^ and GFP^+^ fibers innervating lung slices (conducting airways stained for E-cadherin) taken from *TRPV1^Flp^P2X2^Cre^-FLTG* (A1-3) and *TRPV1^Flp^Tac1^Cre^-FLTG* (B1-3) mice. tdTomato^+^ and GFP^+^ fibers were traced manually, conducting airways (CA) denoted by black lines, blood vessels (BV) denoted by grey lines. Scale bars denote 100μm. C-E, quantification of lung structures innervated by tdTomato^+^ or GFP^+^ fibers in *TRPV1^Flp^P2X2^Cre^-FLTG* (n = 5 animals, 156 CA and 90 BV) and *TRPV1^Flp^Tac1^Cre^-FLTG* (n = 6 animals, 211 CA and 191 BV) mice. C, percentage of conducting airways innervated by reporter-expressing fibers. D, percentage of conducting airways with reporter-expressing fibers that project out to the alveolar region. E, percentage of blood vessels with reporter-expressing fibers.

To specifically map vagal afferent innervation of the lung, we performed vagal microinjection with intersectional mCherry-expressing AAV9. Following injection of homozygous *TRPV1^Flp^P2X2^Cre^*mice with AAV9-ConFon-mCherry (n=3), we found many mCherry^+^ fibers innervating the conducting airways and projecting out into the alveolar region (Fig. 12A). Innervation of some blood vessels was also noted. Following injection of homozygous *TRPV1^Flp^Tac1^Cre^* mice with AAV9-ConFon-mCherry (n=3), we found many mCherry^+^ fibers innervating the conducting airways but none projected out into the alveolar space (Fig. 12B). Some blood vessels were also innervated by mCherry^+^ fibers. Lastly, we only observed mCherry^+^ fibers innervating the largest conducting airways following injection of homozygous *TRPV1^Flp^Tac1^Cre^* mice with AAV9-ConFoff-mCherry (n=3, Fig. 12C).

**Figure 12:**
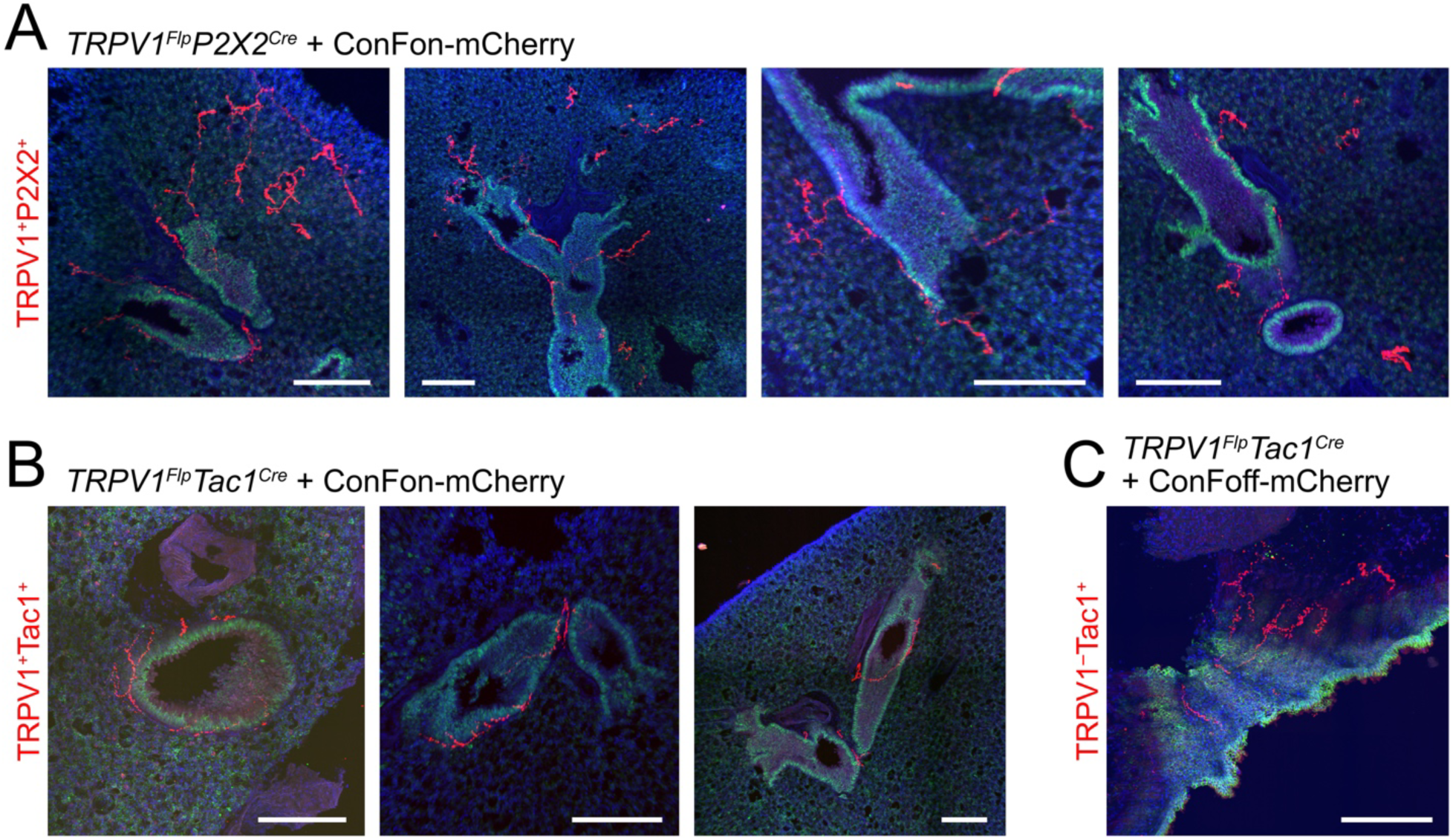
Distinct innervation of the lung by vagal afferent subsets. Representative lung slices showing expression of mCherry (in red) following vagal injection of AAV9-ConFon-mCherry into *TRPV1^Flp^P2X2^Cre^*mice (A) and *TRPV1^Flp^Tac1^Cre^* mice (B), and AAV9-ConFoff-mCherry into *TRPV1^Flp^Tac1^Cre^* mice (C). E-cadherin (in green) and DAPI staining (in blue) also shown. Scale bars denote 200μm.

Overall, our mapping indicates that the lung receives significant innervation by both vagal TRPV1^+^P2X2^+^ afferents and vagal TRPV1^+^Tac1^+^ afferents. While both innervate the conducting airways and some blood vessels, only TRPV1^+^P2X2^+^ afferents project into the alveolar region. Lung innervation by vagal TRPV1^−^Tac1^+^ afferents is limited to the large conducting airways.

### Mapping of tracheal afferent subsets

Confocal imaging of wholemount trachea from *TRPV1^Flp^P2X2^Cre^-FLTG* mice (n=3) revealed very few GFP^+^ fibers but many tdT^+^ fibers in the submucosa of the trachealis and between the hyaline cartilage rings (Fig. 13A). Some tdT^+^ fibers merged into the epithelial layer (not shown). We also found tdT^+^ fibers occasionally innervating the muscle layer of the trachealis. Both GFP^+^ and tdT^+^ fibers were found in nerve tracts in the adventitia (not shown). In *TRPV1^Flp^Tac1^Cre^-FLTG* mice (n=3) we found extensive GFP^+^ innervation of the submucosa of the trachealis and between the rings (Fig. 13B). Some GFP^+^ fibers merged with the epithelium, and some GFP^+^ fibers innervated the trachealis muscle. The very sparse tdT^+^ innervation was found in the submucosa – typically in the trachealis rather than between the cartilage rings. Both GFP^+^ and tdT^+^ fibers were found in nerve tracts in the adventitia (not shown). These data indicate that the vast majority of TRPV1^+^ afferent innervation of the trachea also expresses Tac1 (Table 4).

**Figure 13:**
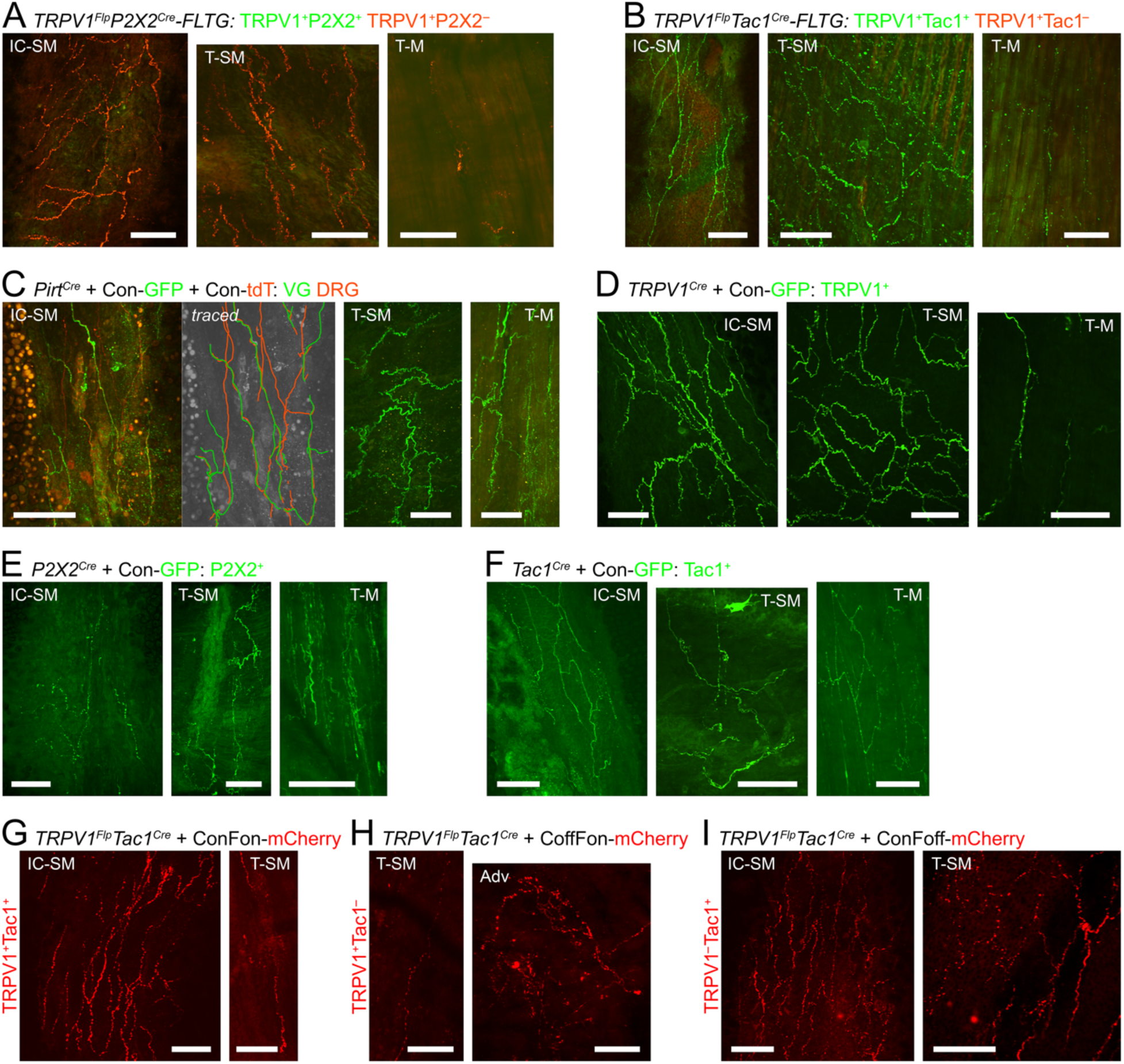
Mapping of sensory afferents innervating the trachea. Confocal images of wholemount trachea longitudinally cut along the ventral midline showing intercartilaginous submucosa (IC-SM), trachealis submucosa (T-SM), trachealis muscle (T-M), or adventitia (Adv) layers. A and B, GFP^+^ and tdTomato^+^ fibers (in green and red, respectively) innervating the trachea of *TRPV1^Flp^P2X2^Cre^-FLTG* (A) and *TRPV1^Flp^Tac1^Cre^-FLTG* (B) mice. C, vagal GFP^+^ fibers (in green) and DRG tdTomato^+^ fibers (in red) innervating the trachea of *Pirt^Cre^* mice following vagal injection with AAV9-Con-GFP and T1-T3 DRG injection with AAV9-Con-tdT. GFP^+^ and tdT^+^ fibers in the IC-SM image were subsequently traced by hand for clarity (*traced*). D-F, GFP^+^ fibers (in green) innervating the trachea of *TRPV1^Cre^*mice (D) or *P2X2^Cre^* mice (E) or *Tac1^Cre^* mice (F) following vagal injection with AAV9-Con-GFP. G-I, mCherry^+^ fibers (in red) innervating the trachea of *TRPV1^Flp^Tac1^Cre^*mice following vagal injection with AAV9-ConFon-mCherry (G) or AAV9-CoffFon-mCherry (H) or AAV9-ConFoff-mCherry (I). For each tissue the left side is rostral and the right side is caudal, except for the T-SM image in A that is rostral towards the top side and caudal towards the bottom side. Scale bars denote 100μm.

**Table 4:** Summary of tracheal innervation by afferent subsets. Scoring: 0 – absent, 1 – very rare, 2 – rare, 3 – occasional, 4 – abundant, 5 – dense.

| Source | Labeled subset | Intercartilaginous |  | Trachealis |  |  |
| --- | --- | --- | --- | --- | --- | --- |
|  |  | Epithelium | Submucosa | Epithelium | Submucosa | Muscle |
| RC-FLTG | TRPV1<sup>+</sup>P2X2<sup>+</sup> | 0 | 2 | 0 | 1 | 0 |
| RC-FLTG | TRPV1<sup>+</sup>P2X2<sup>-</sup> | 2 | 5 | 2 | 5 | 3 |
| RC-FLTG | TRPV1<sup>+</sup>Tac1<sup>+</sup> | 2 | 5 | 2 | 5 | 3 |
| RC-FLTG | TRPV1<sup>+</sup>Tac1<sup>-</sup> | 0 | 1 | 1 | 1 | 0 |
| VG injection | all VG neurons | 2 | 5 | 2 | 5 | 5 |
| DRG injection | all DRG neurons | 0 | 4 | 2 | 2 | 2 |
| VG injection | TRPV1<sup>+</sup> | 1 | 5 | 2 | 4 | 3 |
| VG injection | P2X2<sup>+</sup> | 0 | 3 | 2 | 4 | 5 |
| VG injection | Tac1<sup>+</sup> | 1 | 5 | 1 | 5 | 3 |
| VG injection | TRPV1<sup>+</sup>Tac1<sup>+</sup> | 2 | 5 | 2 | 4 | 3 |
| VG injection | TRPV1<sup>+</sup>Tac1<sup>-</sup> | 0 | 0 | 0 | 2 | 1 |
| VG injection | TRPV1<sup>-</sup>Tac1<sup>+</sup> | 0 | 5 | 0 | 4 | 1 |

Tracheal afferent innervation has been shown to be derived from both vagal and DRG sources ^41,42^. To discriminate between these afferent subsets, we first unilaterally injected AAV9-Con-GFP into the vagal ganglia and AAV9-Con-tdTomato into the thoracic DRG of *Pirt^Cre^*mice (n=3). We found extensive vagal GFP^+^ innervation of the submucosa of the trachealis and intercartilaginous regions (Fig. 13C), some of which projected to the epithelium (not shown). We also identified many vagal GFP^+^ fibers innervating the trachealis muscle layer, which often ran in parallel with the muscle. We found dense and parallel DRG tdT^+^ innervation of the submucosa between the cartilage rings but only sparse innervation of the trachealis (submucosa and muscle) (Fig. 13C). Both vagal and DRG afferent fibers were found in nerve tracts in the adventitia (not shown). Surprisingly, there was little evidence that innervation by either vagal or DRG afferents was greater in the ipsilateral part of the trachea. Following unilateral vagal ganglia injection of *TRPV1^Cre^* mice with AAV9-Con-GFP (n=4), we found extensive GFP^+^ (vagal TRPV1^+^) fibers innervating the submucosa throughout the trachea (sometimes merging with the epithelium), and occasional innervation of the trachealis muscle (Fig. 13D). We found many GFP^+^ fibers in the adventitia (not shown). Following unilateral vagal ganglia injection of *P2X2^Cre^* mice with AAV9-Con-GFP (n=3), we found abundant GFP^+^ (vagal P2X2^+^) fibers innervating the submucosa of the trachealis (sometimes projecting to the epithelium) and less dense GFP^+^ innervation of the intercartilaginous submucosa (Fig. 13E). We also found dense and often parallel GFP^+^ fibers innervating the muscle layer. GFP^+^ fibers were found throughout the adventitia (not shown). Following unilateral vagal ganglia injection of *Tac1^Cre^* mice with AAV9-Con-GFP (n=3), we found extensive GFP^+^ (vagal Tac1^+^) fibers innervating the tracheal submucosa (Fig. 13F). We also identified occasional GFP^+^ fibers innervating the trachealis muscle layer, which commonly ran in parallel with the muscle. GFP^+^ fibers were found throughout the adventitia (not shown). These data indicate that there is extensive innervation of the trachea by vagal afferents that express (albeit to a varying extent) TRPV1, P2X2 and Tac1 (Table 4).

To discriminate between these vagal afferent subsets innervating the trachea, we performed vagal microinjection of *TRPV1^Flp^Tac1^Cre^*mice with intersectional mCherry-expressing AAV9. Injection of AAV9-ConFon-mCherry (n=3) revealed many mCherry^+^ (vagal TRPV1^+^Tac1^+^) fibers innervating the submucosa throughout the trachea, which occasionally projected to the epithelium (Fig. 13G). We found only a few mCherry^+^ fibers innervating the trachealis muscle layer (not shown). Injection of AAV9-CoffFon-mCherry (n=3) labeled only a few mCherry^+^ (vagal TRPV1^+^Tac1^−^) fibers in the trachealis (Fig. 13H), which were typically in the submucosa, but failed to label submucosal fibers between the cartilage rings (not shown). Lastly, AAV9-ConFoff-mCherry injection (n=3) induced mCherry expression in many (vagal TRPV1^−^Tac1^+^) fibers innervating the submucosa throughout the trachea (Fig. 13I). Very few mCherry^+^ fibers innervated the trachealis muscle (not shown). These data demonstrate dense and widespread innervation of the trachea by vagal TRPV1^+^Tac1^+^ and TRPV1^−^Tac1^+^ fibers, with very limited innervation by vagal TRPV1^+^Tac1^−^ fibers (Table 4). Nevertheless, we found abundant fibers from all three subsets in nerve tracts in the adventitia (not shown).

### Chemogenetic activation of cardiopulmonary afferent regulates heart rate and breathing rate

Activation of cardiopulmonary nociceptors evokes robust defensive reflexes including changes in breathing rate and heart rate ^11,12^. Using chemogenetics, we investigated the reflexes evoked by selective stimulation of specific afferent subsets by intravenous stimuli, injected (50μl) via the jugular vein in mice anesthetized with ketamine and dexmedetomidine. We detected the cardiac cycle with ECG electrodes and measured the interval between R waves (RRi). Using diaphragmatic EMG electrodes to detect inspiration, we measured the interval between breaths (BBi). In C57BL/6J control mice (n=7), injection of vehicle or the DREADD receptor agonist clozapine-n-oxide (CNO 300μg/ml) caused little or no effect on RRi or BBi (p>0.05), indicating that CNO is largely inert in mice lacking DREADD receptors in this timeframe (Fig. 14A-D). Subsequent injection of the TRPV1 agonist capsaicin (3μg/ml) evoked an acute bradycardia (increased ΔRRi) and bradypnea (increased ΔBBi), lasting ∼30s (Fig. 14E-H). In 4 out of 7 animals the initial capsaicin-evoked bradypnea was followed by rapid shallow breathing (mean duration of 17.2 ± 10.2s, not shown). To selectively express hM3Dq in vagal TRPV1^+^ neurons, we bilaterally microinjected the vagal ganglia of homozygous *TRPV1^Cre^*mice with AAV9-Con-hM3Dq. In these mice (n=9), 10μg/ml CNO evoked significant bradycardia (p<0.05) and, in some mice, bradypnea, although this did not reach significance (p>0.05)(Fig. 14I-L). Subsequent administration of 100μg/ml CNO evoked significant bradypnea in these mice (p<0.05)(Fig. 14M).

**Figure 14:**
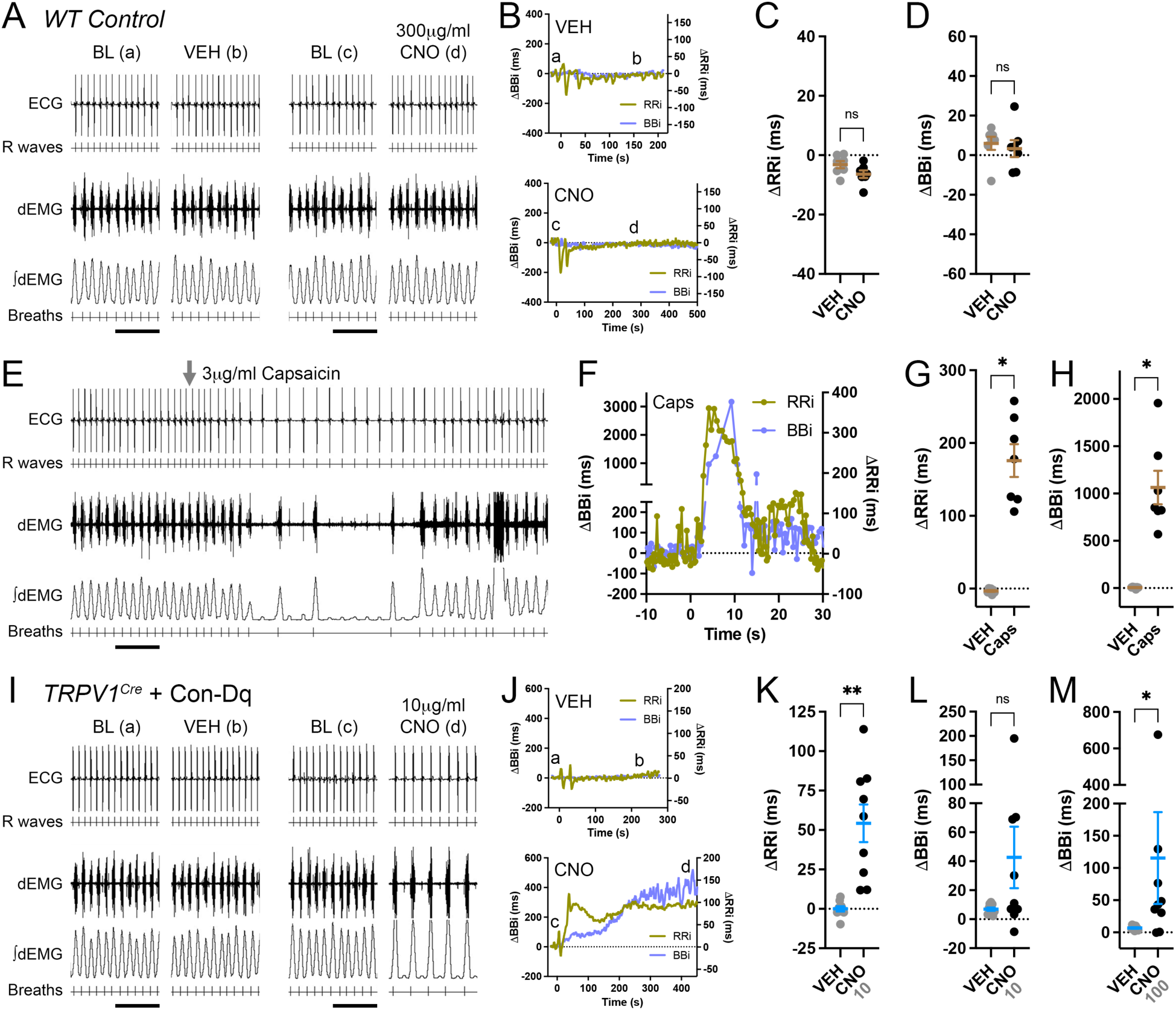
Cardiopulmonary reflexes evoked by intravenous stimulation of TRPV1^+^ bronchopulmonary afferents. Ketamine and dexmedetomidine anesthetized mice with ECG and dEMG leads to measure the ΔRRi and ΔBBi from the pre-stimulus baseline (BL). A-D, wild-type mice treated with i.v. vehicle (VEH) and 300μg/ml CNO (n=7). A, representative ECG with detected R waves and dEMG with integrated dEMG and detected breaths. B, representative rolling average change in RRi (violet line) and BBi (olive line) evoked by vehicle (top) and CNO (bottom), with event markers (a, b, c, d) for the raw traces shown in A (a, b, c, d). C, mean ± SEM ΔRRi. D, mean ± SEM ΔBBi. E-H, wild-type mice treated with i.v. capsaicin (3μg/ml)(n=7). E, representative ECG with detected R waves and dEMG with integrated dEMG and detected breaths. F, representative change in RRi (violet line) and BBi (olive line) evoked by capsaicin. G, mean ± SEM ΔRRi. H, mean ± SEM ΔBBi. I-L, *TRPV1^Cre^* mice following vagal ganglia injection with AAV9-Con-hM3Dq treated with i.v. vehicle and 10μg/ml CNO (n=9). I, representative ECG with detected R waves and dEMG with integrated dEMG and detected breaths. J, representative rolling average change in RRi (violet line) and BBi (olive line) evoked by vehicle (top) and CNO (bottom), with event markers (a, b, c, d) for the raw traces shown in I. K, mean ± SEM ΔRRi for 10μg/ml CNO. L, mean ± SEM ΔBBi for 10μg/ml CNO. M, mean ± SEM ΔBBi for *TRPV1^Cre^* mice following vagal ganglia injection with AAV9-Con-hM3Dq treated with i.v. vehicle and 100μg/ml CNO. Scale bars represent 2s. * represent a significant difference between vehicle and stimulus (p<0.05). ns represent no significant difference between vehicle and stimulus (p>0.05).

In heterozygous *P2X2^Cre^-hM3Dq* mice (n=7), which express the activating hM3Dq DREADD receptor in all P2X2^+^ cells, 3μg/ml CNO evoked robust bradycardia and bradypnea (p<0.05)(Fig. 15A-D). To target vagal TRPV1^+^P2X2^+^ afferents, we bilaterally microinjected the vagal ganglia of homozygous *TRPV1^Flp^P2X2^Cre^* mice with AAV9-ConFon-hM3Dq. In these mice (n=9), 10μg/ml CNO evoked significant bradycardia (p<0.05) and bradypnea (p<0.05)(Fig.15E-H).

**Figure 15:**
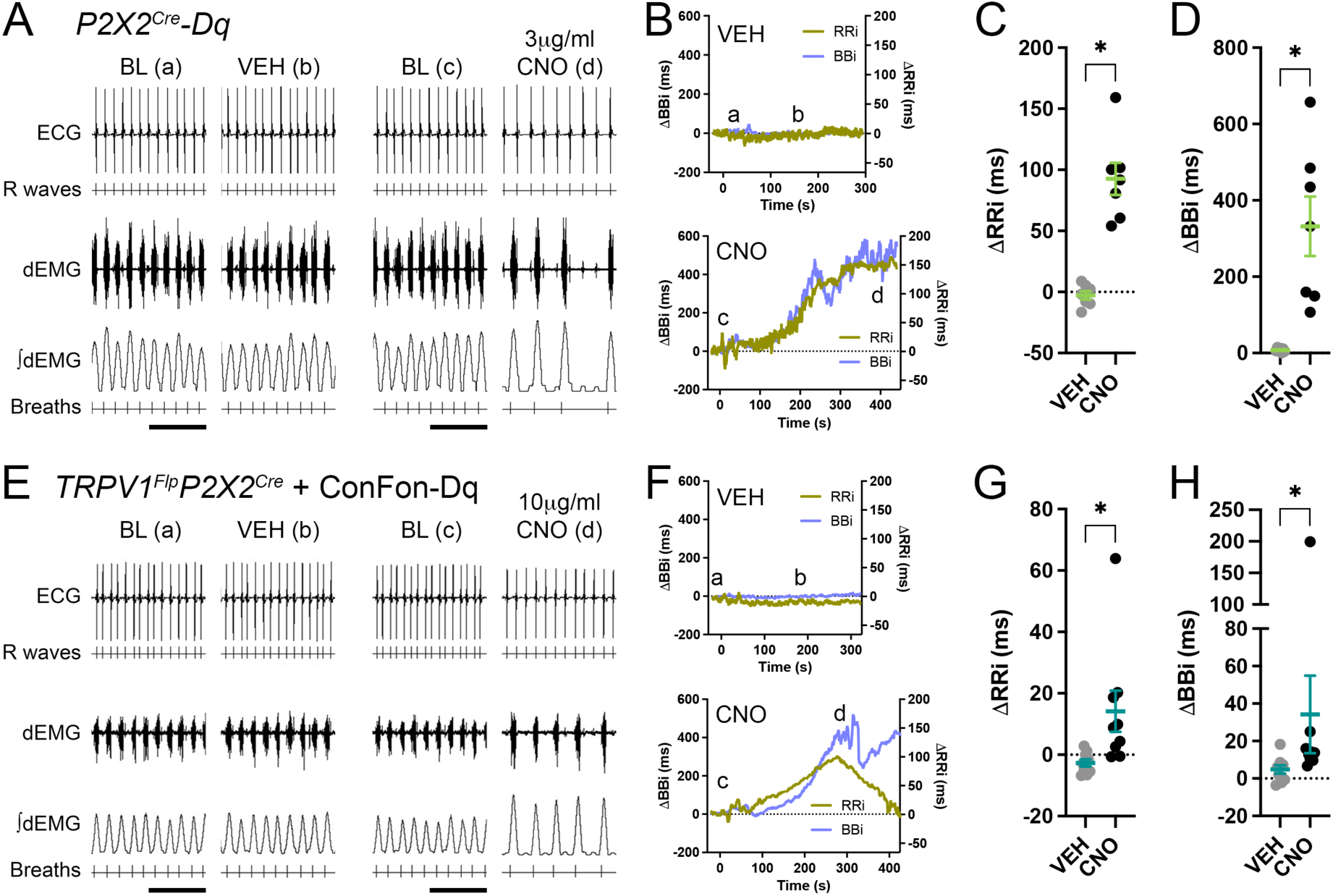
Cardiopulmonary reflexes evoked by intravenous stimulation of P2X2^+^ bronchopulmonary afferents. Ketamine and dexmedetomidine anesthetized mice with ECG and dEMG leads to measure the ΔRRi and ΔBBi from the pre-stimulus baseline (BL). A-D, *P2X2^Cre^-hM3Dq* mice treated with i.v. vehicle and 3μg/ml CNO (n=7). E-H, *TRPV1^Flp^P2X2^Cre^* mice following vagal ganglia injection with AAV9-ConFon-hM3Dq treated with i.v. vehicle and 10μg/ml CNO (n=9). A and E, representative ECG with detected R waves and dEMG with integrated dEMG and detected breaths. B and F, representative rolling average change in RRi (violet line) and BBi (olive line) evoked by vehicle (top) and CNO (bottom), with event markers (a, b, c, d) for the raw traces shown in A or E, respectively. C and G, mean ± SEM ΔRRi. D and H, mean ± SEM ΔBBi. Scale bars represent 2s. * represent a significant difference between vehicle and CNO (p<0.05).

In heterozygous *Tac1^Cre^-hM3Dq* mice (n=6), which express hM3Dq in all Tac1^+^ cells, 3μg/ml CNO evoked robust tachycardia (decreased ΔRRi) and tachypnea (decreased ΔBBi)(p<0.05, Fig. 16A-D). To selectively express hM3Dq in vagal Tac1^+^ neurons, we bilaterally microinjected the vagal ganglia of homozygous *Tac1^Cre^* mice with AAV9-Con-hM3Dq. In these mice (n=6), 10μg/ml CNO evoked tachycardia (p<0.05) but had no effect on breathing rate (p>0.05)(Fig. 16E-H). To target vagal TRPV1^+^Tac1^+^ afferents, we bilaterally microinjected the vagal ganglia of homozygous *TRPV1^Flp^Tac1^Cre^* mice with AAV9-ConFon-hM3Dq. In these mice (n=10), 10μg/ml CNO evoked significant bradycardia (p<0.05), but had little effect on breathing rate (p>0.05)(Fig. 16I-L). Nevertheless, subsequent administration of 100μg/ml CNO evoked significant bradypnea in these mice (p<0.05)(Fig. 16M). Lastly we targeted vagal TRPV1^−^Tac1^+^ afferents by bilateral microinjection of the vagal ganglia of homozygous *TRPV1^Flp^Tac1^Cre^*mice with AAV9-ConFoff-hM3Dq. In these mice (n=6), 10μg/ml CNO evoked tachycardia (p<0.05) but had no effect on breathing rate (p>0.05)(Fig. 16N-Q).

**Figure 16:**
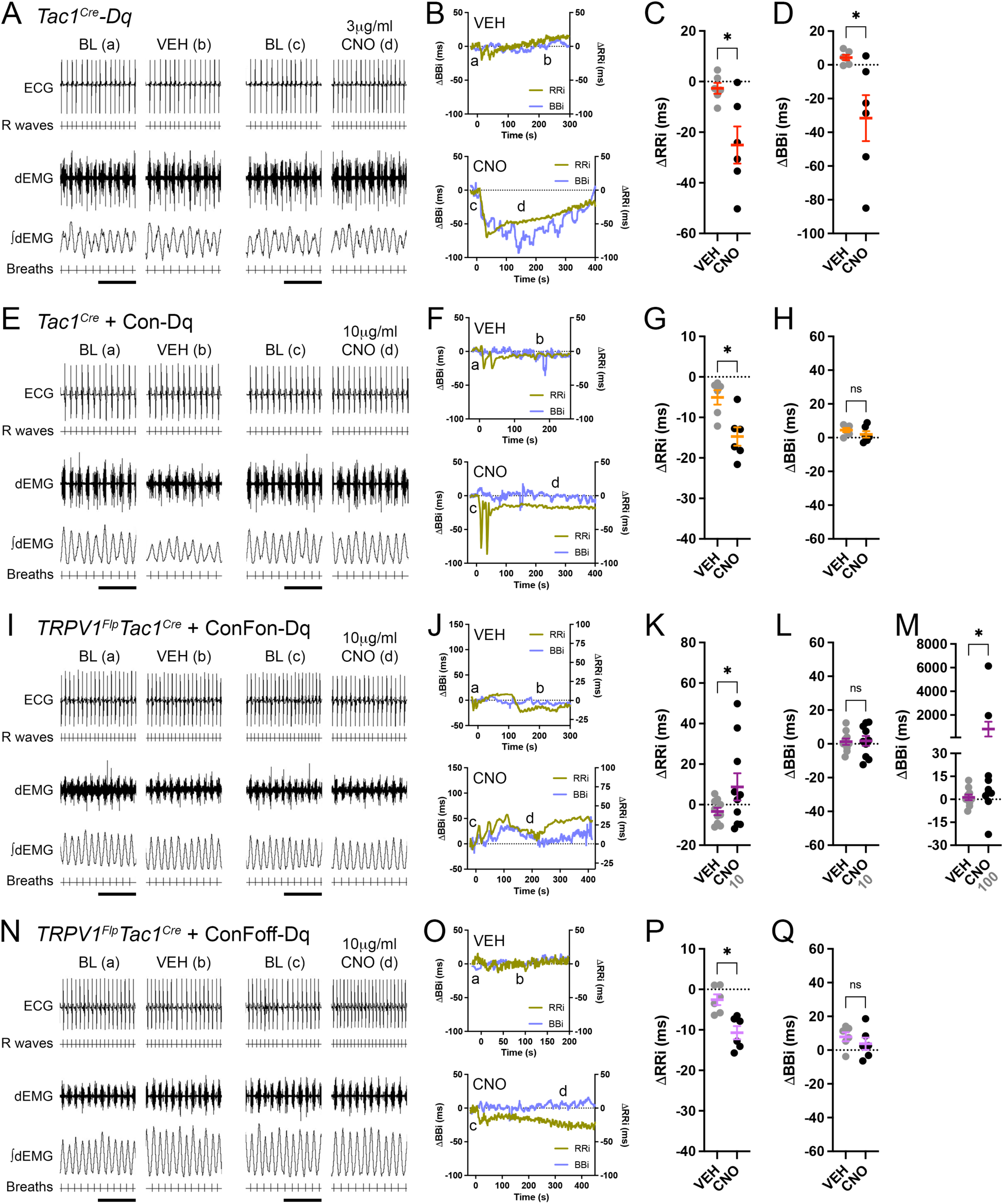
Cardiopulmonary reflexes evoked by intravenous stimulation of Tac1^+^ bronchopulmonary afferents. Ketamine and dexmedetomidine anesthetized mice with ECG and dEMG leads to measure the ΔRRi and ΔBBi from the pre-stimulus baseline (BL). A-D, *Tac1^Cre^-hM3Dq* mice treated with i.v. vehicle and 3μg/ml CNO (n=6). E-H, *Tac1^Cre^* mice following vagal ganglia injection with AAV9-Con-hM3Dq treated with i.v. vehicle and 10μg/ml CNO (n=6). I-L, *TRPV1^Flp^Tac1^Cre^* mice following vagal ganglia injection with AAV9-ConFon-hM3Dq treated with i.v. vehicle and 10μg/ml CNO (n=10). N-Q, *TRPV1^Flp^Tac1^Cre^* mice following vagal ganglia injection with AAV9-ConFoff-hM3Dq treated with i.v. vehicle and 10μg/ml CNO (n=6). A, E, I and N, representative ECG with detected R waves and dEMG with integrated dEMG and detected breaths. B, F, J and O, representative rolling average change in RRi (violet line) and BBi (olive line) evoked by vehicle (top) and CNO (bottom), with event markers (a, b, c, d) for the raw traces shown in A, E, I or N, respectively. C, G, K and P, mean ± SEM ΔRRi. D, H, L and Q, mean ± SEM ΔBBi. M, mean ± SEM ΔBBi for *TRPV1^Flp^Tac1^Cre^* mice following vagal ganglia injection with AAV9-ConFon-hM3Dq treated with i.v. vehicle and 100μg/ml CNO. Scale bars represent 2s. * represent a significant difference between vehicle and stimulus (p<0.05). ns represent no significant difference between vehicle and stimulus (p>0.05).

## Discussion

TRPV1 is a heat- and acid-sensitive ion channel expressed by many nociceptive afferents ^37^, including many innervating the trachea and lung ^43,44^. TRPV1 is essential for nociceptive reflexes evoked by the irritant capsaicin ^45,46^, and is the downstream target of multiple inflammatory pathways ^47^. TRPV1^+^ vagal afferents have been shown to be critical for irritant-evoked bronchospasm ^48–50^, cough ^51^, airway hyperreactivity ^13–16^, pulmonary dysfunction in pneumonia ^17^ and pollution-evoked arrythmia ^52,53^.

Vagal afferent neurons are derived from two distinct embryological sources: placodes-derived neurons in the nodose ganglion, and neural crest-derived neurons in the jugular ganglion ^54^. There is evidence that nodose and jugular afferents differentially regulate cough reflexes ^55^: activation of jugular TRPV1^+^ fibers evoked cough in conscious guinea pigs and augmented cough evoked by citric acid in anesthetized guinea pigs, whereas activation of nodose TRPV1^+^ fibers failed to evoke cough in conscious guinea pigs and inhibited citric acid-evoked cough in anesthetized guinea pigs ^20,56^. Transcriptomics studies have identified substantial phenotypic differences between nodose and jugular neurons ^3,35,39,57^ but little is known about their specific innervation patterns or their contribution to responses other than cough ^23^. This is in part because no single gene or protein is a selective marker of specific afferent subsets. Here, we have used an intersectional approach to label, map and stimulate specific vagal subsets.

TRPV1^+^ vagal afferents have been previously mapped using a *TRPV1^Cre^* mouse strain ^7,10^. Here we developed a *TRPV1^Flp^*mouse strain to use for intersectional genetic manipulation. Reporter expression driven by the *TRPV1^Flp^* allele was highly selective for small-diameter, capsaicin-sensitive afferents. We found little evidence of reporter expression driven by transient TRPV1 expression earlier in development, which has been shown in the *TRPV1^Cre^* strain ^7,58,59^. However, the presence of capsaicin-sensitive reporter-negative neurons suggests the *TRPV1^Flp^* allele did not induce 100% efficient Flp-mediated recombination. This appeared to be more noticeable for the *R26^FLTG/+^*mice than the *R26^ai65f/+^* mice, but could be partially rescued by using homozygous *TRPV1^Flp^* mice.

The purinergic P2X2 is expressed by >80% of nodose afferents but <2% of jugular neurons and ∼10% of DRG neurons ^3,27,35,38,39^. While P2X3 is expressed by almost all vagal afferents, the inclusion of P2X2 in the P2X2/3 heteromer is essential for the selective activation of nodose fibers innervating the airways by αβmATP ^38,60^. Here we used *TRPV1^Flp^P2X2^Cre^* mice crossed with the dual reporter *R26^FLTG^* or vagally injected with AAV9-ConFon-mCherry to differentiate TRPV1^+^ neurons based upon their P2X2 expression. TRPV1^+^P2X2^+^ neurons were found exclusively in the nodose portion of the fused nodose-jugular complex. >90% of the reporter-identified neurons expressed transcript for P2X2 and TRPV1, and were activated by both αβmATP and capsaicin. As expected many tdT^+^ neurons in the *TRPV1^Flp^P2X2^Cre^-FLTG* mice (hypothesized to be TRPV1^+^P2X2^−^) were located in the jugular portion of the ganglion, and the majority lacked P2X2 transcript expression and sensitivity to αβmATP. Nevertheless, some nodose neurons were tdT^+^, and some exhibited P2X2 transcript expression or αβmATP sensitivity suggesting <100% efficiency in the P2X2^Cre^-mediated recombination. The mixed nature of this tdT^+^ group argues against further detailed interpretation.

The gene Tac1 encodes preprotachykinin, the precursor to the tachykinin neuropeptide substance P. Tac1 is expressed in multiple vagal afferent subsets including many TRPV1^+^ and TRPV1^−^ jugular afferents and some TRPV1^−^ nodose afferents ^3,7^. Substance P can be released following afferent activation peripherally where it causes neurogenic bronchospasm ^61^, and centrally where it contributes to synaptic transmission ^49,62,63^. Here we used *TRPV1^Flp^Tac1^Cre^* mice crossed with the dual reporter *R26^FLTG^* or vagally injected with AAV9-ConFon-mCherry to differentiate TRPV1^+^ neurons based upon their Tac1 expression. >90% of TRPV1^+^Tac1^+^ neurons were found in the jugular portion of the nodose-jugular complex. All of the reporter-identified neurons expressed transcript for preprotachykinin and TRPV1, and 90% were activated by capsaicin but not αβmATP. The few TRPV1^+^Tac1^+^ neurons found within the nodose/caudal portion of the complex is consistent with mapping of neural crest-derived Wnt1^+^ neurons ^35^ – suggesting a small number of this population migrate caudally of the jugular ganglion in the mouse. >97% of tdT^+^ neurons in the *TRPV1^Flp^Tac1^Cre^-FLTG* mice (hypothesized to be TRPV1^+^Tac1^−^) were located in the nodose portion of the ganglion, with only 12% expressing Tac1 transcript. This suggests that the recombination of the *R26^FLTG^* allele by *Tac1^Cre^*was more efficient than that by *P2X2^Cre^*.

Vagal injection of *TRPV1^Flp^Tac1^Cre^* mice with AAV9-CoffFon-mCherry identified TRPV1^+^Tac1^−^ neurons. Many were found in the nodose ganglion and these were presumably also P2X2^+^. Nevertheless, this AAV approach found that 15% of TRPV1^+^Tac1^−^ neurons resided in the jugular ganglion. Previous studies in guinea pigs have identified capsaicin-sensitive neurofilament^+^ jugular neurons (i.e. Aδ fibers) that innervate the trachea and lungs but which lack substance P expression ^43^. It should be noted that jugular TRPV1^+^Tac1^−^ neurons were very rarely observed (2.7%) in the *TRPV1^Flp^Tac1^Cre^-FLTG* mice. It is possible that jugular neurons that are TRPV1^+^Tac1^−^ in the adult mouse (when all our studies were performed) transiently expressed Tac1 during development, thus permanently recombining the *R26^FLTG^*allele. Vagal injection of *TRPV1^Flp^Tac1^Cre^* mice with AAV9-ConFoff-mCherry identified TRPV1^−^Tac1^+^ neurons. This large population included both nodose and jugular neurons, although little is known of their current identity. Kupari et al (2019) identified high Tac1 expression in a TRPV1^−^ subset of nodose neurons (NG5), which expressed receptors for inflammatory mediators suggesting a chemosensitive function ^3^.

In the medulla, vagal TRPV1^+^P2X2^+^ afferents terminated in the nTS and area postrema – no terminations in the Pa5 were noted, consistent with previous reports of P2X2^+^ and Phox2b^+^ afferents ^27,28^. TRPV1^+^P2X2^+^ afferents innervated the medial and dorsal subnuclei of the nTS such as SolC, SolG and SolDL, similar to nTS terminations by TRPV1^+^ afferents ^7^. Whereas vagal TRPV1^+^Tac1^+^ afferents exclusively terminated in the Pa5 – we found no evidence that these fibers terminated in the nTS or area postrema. Previous mapping of vagal Tac1^+^ afferents found terminations within the Pa5 and the nTS but little in the area postrema ^7,9^. Our data indicates that only Tac1^+^ vagal afferents lacking TRPV1 expression project to the nTS. Indeed TRPV1^−^Tac1^+^ afferents terminate preferentially in lateral and ventral subnuclei (e.g. SolVL), distinct from TRPV1^+^ terminations. It should be noted that SolC microinjection of NK antagonists in rats inhibited the bradypnea evoked by SolC microinjection of capsaicin ^64^, which was taken as evidence of capsaicin-sensitive jugular afferent (i.e. TRPV1^+^Tac1^+^) innervation of the nTS. However, such functional data does not explicitly identify a role for *afferent* substance P, and evidence from Tac1 reporter mice ^7^ and single neuron transcriptomics ^65^ suggest many nTS neurons themselves express substance P. Lastly, we found TRPV1^+^Tac1^−^ afferents terminated in the nTS, area postrema and the Pa5. As described above, this group represents two distinct subsets: nodose TRPV1^+^P2X2^+^Tac1^−^ and jugular TRPV1^+^P2X2^−^Tac1^−^ afferents. The Pa5 terminations are derived exclusively from the latter group that may be capsaicin-sensitive jugular Aδ fibers. It is not currently known if some TRPV1^+^P2X2^−^Tac1^−^ afferents also project to the nTS and area postrema.

In the lung, vagal TRPV1^+^P2X2^+^ afferents innervated most conducting airways and a small number of pulmonary blood vessels. Strikingly, TRPV1^+^P2X2^+^ afferents also projected from many conducting airways into the alveolar region. These data are similar to the innervation of the lung by vagal TRPV1^+^ afferents and nodose 5HT3^+^ afferents ^10^. TRPV1^+^Tac1^+^ afferents also innervated most conducting airways and a few pulmonary blood vessels. TRPV1^+^Tac1^+^ afferents labeled by anterograde tracing from the vagal ganglion yielded identical innervation patterns as the total TRPV1^+^Tac1^+^ population, indicating jugular neurons as the primary source of this afferent subset. In stark contrast to TRPV1^+^P2X2^+^ afferents, no TRPV1^+^Tac1^+^ afferents projected from the conducting airways into the alveolar region, consistent with previous mapping of the lung by vagal Tac1^+^ afferents ^9,10^. We also found TRPV1^−^Tac1^+^ afferents innervating the large conducting airways, although these were infrequent.

In contrast to the lung, we found very little evidence of TRPV1^+^P2X2^+^ afferents innervating the trachea (other than the adventitia), and these rare fibers tended to innervate the submucosa between the cartilage rings. Nevertheless, we observed abundant vagal P2X2^+^ innervation of the submucosa and muscle layers of the trachealis and occasional innervation of the intercartilaginous submucosa, consistent with previous nodose afferent mapping studies with P2X2- and Phox2b-reporter mice ^27,28^. We therefore conclude that the majority of nodose afferents innervating the trachea lack TRPV1 expression, consistent with widespread insensitivity to capsaicin in recordings of nodose tracheal afferents ^43,66^. Instead, we found dense innervation of the tracheal submucosa by TRPV1^+^Tac1^+^ afferents, which occasionally projected into the epithelial layer, which is consistent with the colocalization of TRPV1- and substance P-immunoreactivity in this tracheal layer ^67^. Multiple independent lines of evidence indicate that these afferents are largely projected from jugular neurons. Firstly, our anterograde tracing showed denser vagal innervation of the trachea compared to DRG innervation, particular in the trachealis, and this is consistent with previous retrograde tracing studies in rats and guinea pigs have shown that <10% of tracheal afferent innervation projected from the DRG ^41,42^. Secondly, anterograde tracing of vagal TRPV1^+^ afferents demonstrated dense innervation of the tracheal submucosa, consistent with previous immunohistochemical studies ^68,69^, which was very similar to the innervation patterns of TRPV1^+^Tac1^+^ afferents here but unlike TRPV1^+^P2X2^+^ afferents which were rarely observed. Thirdly, anterograde tracing of vagal TRPV1^+^Tac1^+^ afferents produced very similar innervation patterns to the total TRPV1^+^Tac1^+^ afferent population. Taken together, our data suggest that much or most of the TRPV1^+^ afferent innervation of the mouse trachea is projected from Tac1^+^ jugular afferents, and this is consistent with single afferent recordings of capsaicin-sensitive jugular afferents innervating the trachea and capsaicin-induced neuropeptide-mediated tracheal contraction studies in guinea pigs ^43,61^. The TRPV1^+^Tac1^+^ afferents innervating the intercartilaginous submucosa were arranged in parallel anterolateral segmental arrays ^70^, and when anterogradely traced by unilateral vagal AAV injections appeared to innervate ipsilateral and contralateral sides of the trachea equally (as did the overall vagal and DRG innervation). Whereas the TRPV1^+^Tac1^+^ afferents innervating the submucosa of the trachealis (and the limited muscle innervation too) were arranged in large radially branching networks ^70^.

The tracheal submucosa was also densely innervated by vagal TRPV1^−^Tac1^+^ afferents, consistent with previous immunohistochemistry studies in guinea pigs ^67^. It is not clear if these fibers, which largely resemble TRPV1^+^Tac1^+^ afferents in terminal structure, are projected from nodose or jugular neurons, but the likelihood that they are the latter is greater in the intercartilaginous submucosa because it has sparser vagal P2X2^+^ innervation than the trachealis submucosa. We consistently found less TRPV1^+^Tac1^+^ and TRPV1^−^Tac1^+^ innervation of the muscle layer compared to the submucosa. Instead muscle fibers, which received only sparse DRG innervation, were densely innervated by parallel P2X2^+^ afferents, which (as discussed above) are highly likely to lack TRPV1 expression. Lastly, anterograde tracing of vagal TRPV1^+^Tac1^−^ innervation revealed some fibers within the trachealis submucosa. Given that TRPV1^+^P2X2^+^ afferents were very rarely observed in this tissue, it is possible these TRPV1^+^Tac1^−^ afferents are capsaicin-sensitive jugular Aδ fibers that have been identified in electrophysiological recordings of guinea pig trachea afferents ^43^.

Our data clearly demonstrating jugular TRPV1^+^Tac1^+^ afferents innervating the trachea and lungs and only projecting to the Pa5 (rather than the nTS) is inconsistent with the lack of labeled jugular neurons or terminations in the Pa5 following AAV injections or instillations into the mouse airways ^7,9,10,28^. As has been argued elsewhere ^10^, there appears to be a AAV tropism effect for the retrograde labeling (but not anterograde labeling) of all neural crest-derived afferents innervating the mouse airways, because airway DRG afferents are also not retrogradely labeled by AAV. Nevertheless, a non-vector retrograde tracer (DiI) labeled jugular neurons projecting to the mouse airways ^35^, providing further evidence that the mouse has similar neural crest-derived innervation patterns to the rat ^40^ and guinea pig ^38,43,44,49^, and possibly humans ^71^.

The activation of nociceptive afferents innervating the airways evokes robust defensive reflexes including cough, dyspnea, hypersecretion, bronchospasm, changes in respiratory drive, heart rate and blood pressure ^11,12^. Here, an i.v. bolus of capsaicin evoked rapid and robust bradycardia and bradypnea/apnea in anesthetized C57BL/6 mice, consistent with the well-characterized cardiopulmonary reflex that is abolished by vagotomy ^11^. In some mice the capsaicin-evoked bradypnea was followed by tachypnea (rapid shallow breathing). As there are no known selective stimuli for specific vagal afferent subsets, we investigated the functional role of genetically defined bronchopulmonary using chemogenetics (hM3Dq DREADD). CNO-mediated activation of vagal TRPV1^+^ afferents evoked bradycardia and bradypnea, mimicking the effect of capsaicin. Interestingly, the evoked bradycardia occurred considerably earlier than the bradypnea, and the latter was only significant at higher CNO doses. This suggests some independence between the TRPV1^+^ pathways in regulating heart rate and breathing rate. No tachypnea was noted, although this may reflect the slower onset and longer duration of the CNO-evoked responses compared to capsaicin. CNO-mediated activation of P2X2^+^ afferents (which are likely nodose afferents due to the limited of P2X2 expression in the DRG and trigeminal ganglion ^27^) and vagal TRPV1^+^P2X2^+^ afferents both evoked bradycardia and bradypnea. These responses are similar to the bradycardia, hypotension and apnea evoked by selective 5HT3 agonists (e.g. phenylbiguanide) administered into the pulmonary circulation of mice ^72^. However, 5HT3 agonist-evoked apnea has been shown to be followed by rapid shallow breathing in rats, guinea pigs and cats ^11,20,73,74^. That mouse nodose afferents can evoke tachypnea has been shown by optogenetic experiments: stimulation of Npy2r^+^ afferents innervating the area postrema evoked apnea followed by rapid shallow breathing ^75^, and low frequency stimulation of Phox2b^+^ afferents in the right cervical vagus (although not the left vagus) evoked tachypnea ^28^. The discrepancy in tachypneic responses between these optogenetic experiments and our data may be due to the selective stimulation of airway afferents here, or because the kinetics or magnitude of the chemogenetic responses were not sufficient to trigger tachypnea.

We found considerable complexity in the reflex response to the activation of Tac1^+^ afferents. Chemogenetic activation of the total cardiopulmonary Tac1^+^ innervation evoked rapid and robust tachycardia and tachypnea. Specific activation of vagal Tac1^+^ afferents, however, evoked tachycardia without changing breathing rate – suggesting the evoked tachypnea was mediated by Tac1^+^ DRG afferents (which innervate the large conducting airways and large pulmonary vessels ^10^), although this was not further tested in the current study. The vagally-mediated Tac1^+^ afferent responses were mimicked by chemogenetic activation of vagal TRPV1^−^Tac1^+^ afferents, whereas CNO-mediated activation of TRPV1^+^Tac1^+^ afferents evoked bradycardia and bradypnea. Thus it appears that the vagal TRPV1^+^Tac1^+^ afferent-mediated responses are overcome/suppressed by vagal TRPV1^−^Tac1^+^ afferent-mediated responses when simultaneously stimulated. While the TRPV1^+^Tac1^+^ afferents involved in these responses are highly likely to be jugular afferents projecting to the Pa5, it is not clear if nodose or jugular pathways mediated the vagal TRPV1^−^Tac1^+^ afferent responses. Our data conflicts with a report that low frequency optogenetic stimulation of jugular Wnt1^+^ afferents in the right cervical vagus (but not the left vagus) evoked tachypnea without changing heart rate ^28^.

Despite the observation that i.v. capsaicin evoked rapid shallow breathing after the initial apnea in some mice, we found no evidence that i.v. CNO-mediated chemogenetic stimulation of specific vagal afferent subsets evoked tachypnea. Although this could implicate the DRG Tac1^+^ in the capsaicin-evoked tachypnea, this is unlikely because many Tac1^+^ fibers in the DRG that innervate the mouse lungs are TRPV1^−^ ^10^, and vagotomy eliminated the vast majority of the reflex evoked by i.v. irritants in other mammalian species ^11,20,73^. Instead, our mouse data is consistent with the hypothesis that irritant-evoked tachypnea is a consequence of the initial apnea ^11^. It is possible that rapid and robust responses to bolus capsaicin are less representative of inflammation- and infection-induced activation of airway nociceptive fibers than the slower DREADD-mediated chemogenetics, and so it is debatable which response actually contributes to reflexes in pathophysiological conditions. However, it should be noted that slow i.v. infusion of capsaicin triggers tachypnea in dogs without apnea ^76,77^, suggesting possible species differences.

Cardiopulmonary reflexes evoked by i.v. irritants have long been thought to be mediated by vagal C fiber afferents specifically innervating the alveoli – the so-called Juxtacapillary receptors or J-receptors ^11^. Our mapping of vagal afferent subsets in the mouse shows that only TRPV1^+^P2X2^+^ afferents (i.e. nodose C-fibers) innervate the alveolar region, and that TRPV1^−^Tac1^+^ and TRPV1^+^Tac1^+^ afferents innervate only the conducting airways and large blood vessels in the lung. Importantly, we observed significant responses to i.v. administration of CNO in mice expressing hM3Dq in TRPV1^−^Tac1^+^ and TRPV1^+^Tac1^+^ afferents. This suggests that, in the mouse, irritants within the pulmonary circulation can sufficiently activate afferent terminals innervating other compartments of the lung.

A limitation of this study was the partial inefficiency of the recombination by Cre and Flp. While selectivity was high (i.e. very few false positives), we found evidence of failures of recombination in some neurons (i.e. false negatives). This is likely dependent on the number of recombinase molecules ^78,79^, with the neurotransmitter Tac1 likely to be expressed at higher levels than the membrane receptor P2X2. While similar efficiencies for Cre and Flp_O_ recombination has been reported in HEK293 cells ^33^, mouse studies have shown more efficient recombination by Cre compared to Flp_O_ ^29,80^. Previous studies suggest that recombinase efficiency may vary across specific afferent subsets ^81,82^, but the extent to which this contributed to the current study is unknown. Decreased efficiency would be further exacerbated by less than 100% transfection by the AAV9 intraganglionic injections. Nevertheless, the approach taken here was sufficient to reproducibly map and modulate distinct vagal afferent populations.

In summary, using an intersectional approach we have determined the innervation patterns and reflex function of multiple nociceptive vagal afferent subsets. We have shown that nodose TRPV1^+^ and jugular TRPV1^+^ afferents have distinct innervation patterns in the lower airways and the medulla. Despite previous data demonstrating opposing effects of these subsets on the cough reflex in guinea pigs ^20,56^, we found they largely evoked similar bradypnea and bradycardia when activated by i.v. stimulants. Our data has identified TRPV1^−^Tac1^+^ afferent subsets (both vagal and non-vagal) that evoke tachypnea and tachycardia when activated by i.v. stimulants. Little is known of the activation profile or physiological role of TRPV1^−^Tac1^+^ afferents innervating the airways, and this should be the focus of future studies. It should be noted that gene expression in vagal afferents is not static and that inflammation and infection can induce *de novo* expression of multiple genes including TRPV1 and Tac1 ^83,84^. How such neuroplasticity impacts vagal afferent innervation and function has yet to be elucidated.

## Acknowledgements

This work was supported by the National Institutes of Health’s Office of Director SPARC Commonfund program (OT2OD023854), the National Institute for Neurological Disorders and Stroke (U01NS113868), the National Heart, Lung and Blood Institute (R01HL152219 and T32HL160529) and the National Institute of Diabetes and Digestive and Kidney Diseases (R01DK141890).

## References

1. Reeh PW, Fischer MJM. Nobel somatosensations and pain. Pflugers Arch 2022; 474(4): 405–20.

2. Usoskin D, Furlan A, Islam S, Abdo H, Lonnerberg P, Lou D, Hjerling-Leffler J, Haeggstrom J, Kharchenko O, Kharchenko PV, Linnarsson S, Ernfors P. Unbiased classification of sensory neuron types by large-scale single-cell RNA sequencing. Nat Neurosci 2015; 18(1): 145–53.

3. Kupari J, Haring M, Agirre E, Castelo-Branco G, Ernfors P. An Atlas of Vagal Sensory Neurons and Their Molecular Specialization. Cell Rep 2019; 27(8): 2508–23.e4.

4. Bassi JK, Connelly AA, Butler AG, Liu Y, Ghanbari A, Farmer DGS, Jenkins MW, Melo MR, McDougall SJ, Allen AM. Analysis of the distribution of vagal afferent projections from different peripheral organs to the nucleus of the solitary tract in rats. J Comp Neurol 2022; 530(17): 3072–103.

5. Ran C, Boettcher JC, Kaye JA, Gallori CE, Liberles SD. A brainstem map for visceral sensations. Nature 2022; 609(7926): 320–6.

6. Prescott SL, Liberles SD. Internal senses of the vagus nerve. Neuron 2022; 110(4): 579–99.

7. Kim SH, Hadley SH, Maddison M, Patil M, Cha BJ, Kollarik M, Taylor-Clark TE. Mapping of sensory nerve subsets within the vagal ganglia and the brainstem using reporter mice for Pirt, TRPV1, 5HT3 and Tac1 expression. eNeuro 2020; 7(2).

8. Lee LY, Yu J. Sensory nerves in lung and airways. Comprehensive Physiology 2014; 4(1): 287–324.

9. Su Y, Barr J, Jaquish A, Xu J, Verheyden JM, Sun X. Identification of Lung Innervating Sensory Neurons and Their Target Specificity. Am J Physiol Lung Cell Mol Physiol 2021; 322(1): 50–63.

10. Kim SH, Patil MJ, Hadley SH, Bahia PK, Butler SG, Madaram M, Taylor-Clark TE. Mapping of the Sensory Innervation of the Mouse Lung by Specific Vagal and Dorsal Root Ganglion Neuronal Subsets. eNeuro 2022; 9(2).

11. Coleridge JC, Coleridge HM. Afferent vagal C fibre innervation of the lungs and airways and its functional significance. Rev Physiol Biochem Pharmacol 1984; 99: 1–110.

12. Mazzone SB, Undem BJ. Vagal Afferent Innervation of the Airways in Health and Disease. Physiol Rev 2016; 96(3): 975–1024.

13. Trankner D, Hahne N, Sugino K, Hoon MA, Zuker C. Population of sensory neurons essential for asthmatic hyperreactivity of inflamed airways. Proc Natl Acad Sci U S A 2014; 111(31): 11515–20.

14. Bessac BF, Sivula M, von Hehn CA, Escalera J, Cohn L, Jordt SE. TRPA1 is a major oxidant sensor in murine airway sensory neurons. J Clin Invest 2008; 118(5): 1899–910.

15. Balestrini A, Joseph V, Dourado M, Reese RM, Shields SD, Rougé L, Bravo DD, Chernov-Rogan T, Austin CD, Chen H, Wang L, Villemure E, Shore DGM, Verma VA, Hu B, Chen Y, Leong L, Bjornson C, Hötzel K, Gogineni A, Lee WP, Suto E, Wu X, Liu J, Zhang J, Gandham V, Wang J, Payandeh J, Ciferri C, Estevez A, Arthur CP, Kortmann J, Wong RL, Heredia JE, Doerr J, Jung M, Vander Heiden JA, Roose-Girma M, Tam L, Barck KH, Carano RAD, Ding HT, Brillantes B, Tam C, Yang X, Gao SS, Ly JQ, Liu L, Chen L, Liederer BM, Lin JH, Magnuson S, Chen J, Hackos DH, Elstrott J, Rohou A, Safina BS, Volgraf M, Bauer RN, Riol-Blanco L. A TRPA1 inhibitor suppresses neurogenic inflammation and airway contraction for asthma treatment. J Exp Med 2021; 218(4).

16. Xing Y, Nho Y, Lawson K, Zhu Y, Ellison AE, Chang MY, Hancock W, Han L. MrgprC11(+) Jugular Neurons Control Airway Hyperresponsiveness in Allergic Airway Inflammation. Am J Respir Cell Mol Biol 2024.

17. Baral P, Umans BD, Li L, Wallrapp A, Bist M, Kirschbaum T, Wei Y, Zhou Y, Kuchroo VK, Burkett PR, Yipp BG, Liberles SD, Chiu IM. Nociceptor sensory neurons suppress neutrophil and γδ T cell responses in bacterial lung infections and lethal pneumonia. Nat Med 2018; 24(4): 417–26.

18. Canning BJ, Undem BJ. Evidence that distinct neural pathways mediate parasympathetic contractions and relaxations of guinea-pig trachealis. J Physiol 1993; 471: 25–40.

19. Muroi Y, Ru F, Chou YL, Carr MJ, Undem BJ, Canning BJ. Selective inhibition of vagal afferent nerve pathways regulating cough using Nav 1.7 shRNA silencing in guinea pig nodose ganglia. Am J Physiol Regul Integr Comp Physiol 2013; 304(11): R1017–23.

20. Chou YL, Mori N, Canning BJ. Opposing effects of bronchopulmonary C-fiber subtypes on cough in guinea pigs. Am J Physiol Regul Integr Comp Physiol 2018; 314(3): R489–r98.

21. Hooper JS, Stanford KR, Alencar PA, Alves NG, Breslin JW, Dean JB, Morris KF, Taylor-Clark TE. Nociceptive pulmonary-cardiac reflexes are altered in the Spontaneously Hypertensive rat. J Physiol 2019; 597(13): 3255–79.

22. Chou YL, Scarupa MD, Mori N, Canning BJ. Differential effects of airway afferent nerve subtypes on cough and respiration in anesthetized guinea pigs. Am J Physiol Regul Integr Comp Physiol 2008; 295(5): R1572–84.

23. Taylor-Clark TE. Molecular identity, anatomy, gene expression and function of neural crest vs. placode-derived nociceptors in the lower airways. Neurosci Lett 2021; 742: 135505.

24. Wang J, Kollarik M, Ru F, Sun H, McNeil B, Dong X, Stephens G, Korolevich S, Brohawn P, Kolbeck R, Undem B. Distinct and common expression of receptors for inflammatory mediators in vagal nodose versus jugular capsaicin-sensitive/TRPV1-positive neurons detected by low input RNA sequencing. PLoS One 2017; 12(10): e0185985.

25. Mazzone SB, Tian L, Moe AAK, Trewella MW, Ritchie ME, McGovern AE. Transcriptional Profiling of Individual Airway Projecting Vagal Sensory Neurons. Mol Neurobiol 2019; 57(2): 949–63.

26. Nonomura K, Woo SH, Chang RB, Gillich A, Qiu Z, Francisco AG, Ranade SS, Liberles SD, Patapoutian A. Piezo2 senses airway stretch and mediates lung inflation-induced apnoea. Nature 2017; 541(7636): 176–81.

27. Kim SH, Bahia PK, Patil M, Sutton S, Sowells I, Hadley SH, Kollarik M, Taylor-Clark TE. Development of a mouse reporter strain for the purinergic P2X(2) receptor. eNeuro 2020; 7(4).

28. Moe AAK, Bautista TG, Trewella MW, Korim WS, Yao ST, Behrens R, Driessen AK, McGovern AE, Mazzone SB. Investigation of vagal sensory neurons in mice using optical vagal stimulation and tracheal neuroanatomy. iScience 2024; 27(3): 109182.

29. Daigle TL, Madisen L, Hage TA, Valley MT, Knoblich U, Larsen RS, Takeno MM, Huang L, Gu H, Larsen R, Mills M, Bosma-Moody A, Siverts LA, Walker M, Graybuck LT, Yao Z, Fong O, Nguyen TN, Garren E, Lenz GH, Chavarha M, Pendergraft J, Harrington J, Hirokawa KE, Harris JA, Nicovich PR, McGraw MJ, Ollerenshaw DR, Smith KA, Baker CA, Ting JT, Sunkin SM, Lecoq J, Lin MZ, Boyden ES, Murphy GJ, da Costa NM, Waters J, Li L, Tasic B, Zeng H. A Suite of Transgenic Driver and Reporter Mouse Lines with Enhanced Brain-Cell-Type Targeting and Functionality. Cell 2018; 174(2): 465–80.e22.

30. Harris JA, Hirokawa KE, Sorensen SA, Gu H, Mills M, Ng LL, Bohn P, Mortrud M, Ouellette B, Kidney J, Smith KA, Dang C, Sunkin S, Bernard A, Oh SW, Madisen L, Zeng H. Anatomical characterization of Cre driver mice for neural circuit mapping and manipulation. Frontiers in neural circuits 2014; 8: 76.

31. Kim YS, Anderson M, Park K, Zheng Q, Agarwal A, Gong C, Saijilafu, Young L, He S, LaVinka PC, Zhou F, Bergles D, Hanani M, Guan Y, Spray DC, Dong X. Coupled Activation of Primary Sensory Neurons Contributes to Chronic Pain. Neuron 2016; 91(5): 1085–96.

32. Cavanaugh DJ, Chesler AT, Jackson AC, Sigal YM, Yamanaka H, Grant R, O’Donnell D, Nicoll RA, Shah NM, Julius D, Basbaum AI. Trpv1 reporter mice reveal highly restricted brain distribution and functional expression in arteriolar smooth muscle cells. J Neurosci 2011; 31(13): 5067–77.

33. Fenno LE, Mattis J, Ramakrishnan C, Hyun M, Lee SY, He M, Tucciarone J, Selimbeyoglu A, Berndt A, Grosenick L, Zalocusky KA, Bernstein H, Swanson H, Perry C, Diester I, Boyce FM, Bass CE, Neve R, Huang ZJ, Deisseroth K. Targeting cells with single vectors using multiple-feature Boolean logic. Nature methods 2014; 11(7): 763–72.

34. Paxinos G, Franklin B. Mouse Brain in Stereotaxic Coordinates. 4th Edition ed: Elsevier; 2012.

35. Nassenstein C, Taylor-Clark TE, Myers AC, Ru F, Nandigama R, Bettner W, Undem BJ. Phenotypic distinctions between neural crest and placodal derived vagal C-fibres in mouse lungs. J Physiol 2010; 588(Pt 23): 4769–83.

36. Nassenstein C, Kwong K, Taylor-Clark T, Kollarik M, Macglashan DM, Braun A, Undem BJ. Expression and function of the ion channel TRPA1 in vagal afferent nerves innervating mouse lungs. J Physiol 2008; 586(6): 1595–604.

37. Caterina MJ, Schumacher MA, Tominaga M, Rosen TA, Levine JD, Julius D. The capsaicin receptor: a heat-activated ion channel in the pain pathway. Nature 1997; 389(6653): 816–24.

38. Kwong K, Kollarik M, Nassenstein C, Ru F, Undem BJ. P2X2 Receptors Differentiate Placodal vs Neural Crest C-fiber Phenotypes Innervating Guinea Pig Lungs and Esophagus. Am J Physiol Lung Cell Mol Physiol 2008; 295(5): L858–65.

39. Kollarik M, Ru F, Undem BJ. Phenotypic distinctions between the nodose and jugular TRPV1-positive vagal sensory neurons in the cynomolgus monkey. Neuroreport 2019; 30(8): 533–7.

40. McGovern AE, Davis-Poynter N, Farrell MJ, Mazzone SB. Transneuronal tracing of airways-related sensory circuitry using herpes simplex virus 1, strain H129. Neuroscience 2012; 207: 148–66.

41. Springall DR, Cadieux A, Oliveira H, Su H, Royston D, Polak JM. Retrograde tracing shows that CGRP-immunoreactive nerves of rat trachea and lung originate from vagal and dorsal root ganglia. J Auton Nerv Syst 1987; 20(2): 155–66.

42. Kummer W, Fischer A, Kurkowski R, Heym C. The sensory and sympathetic innervation of guinea-pig lung and trachea as studied by retrograde neuronal tracing and double-labelling immunohistochemistry. Neuroscience 1992; 49(3): 715–37.

43. Ricco MM, Kummer W, Biglari B, Myers AC, Undem BJ. Interganglionic segregation of distinct vagal afferent fibre phenotypes in guinea-pig airways. J Physiol 1996; 496 ( Pt 2): 521–30.

44. Undem BJ, Chuaychoo B, Lee MG, Weinreich D, Myers AC, Kollarik M. Subtypes of vagal afferent C-fibres in guinea-pig lungs. J Physiol 2004; 556(Pt 3): 905–17.

45. Caterina MJ, Leffler A, Malmberg AB, Martin WJ, Trafton J, Petersen-Zeitz KR, Koltzenburg M, Basbaum AI, Julius D. Impaired nociception and pain sensation in mice lacking the capsaicin receptor. Science 2000; 288(5464): 306–13.

46. Kollarik M, Undem BJ. Activation of bronchopulmonary vagal afferent nerves with bradykinin, acid and vanilloid receptor agonists in wild-type and TRPV1-/- mice. J Physiol 2004; 555(Pt 1): 115–23.

47. Rosenbaum T, Simon SA. TRPV1 Receptors and Signal Transduction. In: Liedtke WB, Heller S, eds. TRP Ion Channel Function in Sensory Transduction and Cellular Signaling Cascades. Boca Raton (FL): CRC Press/Taylor & Francis Copyright © 2007, Taylor & Francis Group, LLC.; 2007.

48. Canning BJ, Reynolds SM, Mazzone SB. Multiple mechanisms of reflex bronchospasm in guinea pigs. J Appl Physiol (1985) 2001; 91(6): 2642–53.

49. Mazzone SB, Canning BJ. Synergistic interactions between airway afferent nerve subtypes mediating reflex bronchospasm in guinea pigs. Am J Physiol Regul Integr Comp Physiol 2002; 283(1): R86–98.

50. Ellis JL, Undem BJ. Non-adrenergic, non-cholinergic contractions in the electrically field stimulated guinea-pig trachea. Br J Pharmacol 1990; 101(4): 875–80.

51. Tanaka M, Maruyama K. Mechanisms of capsaicin- and citric-acid-induced cough reflexes in guinea pigs. J Pharmacol Sci 2005; 99(1): 77–82.

52. Ghelfi E, Rhoden CR, Wellenius GA, Lawrence J, Gonzalez-Flecha B. Cardiac oxidative stress and electrophysiological changes in rats exposed to concentrated ambient particles are mediated by TRP-dependent pulmonary reflexes. Toxicol Sci 2008; 102(2): 328–36.

53. Taylor-Clark TE. Air Pollution-Induced Autonomic Modulation. Physiology (Bethesda) 2020; 35(6): 363–74.

54. Baker CV, Bronner-Fraser M. Vertebrate cranial placodes I. Embryonic induction. Dev Biol 2001; 232(1): 1–61.

55. Taylor-Clark TE. Peripheral neural circuitry in cough. Curr Opin Pharmacol 2015; 22: 9–17.

56. Mazzone SB, Mori N, Canning BJ. Synergistic interactions between airway afferent nerve subtypes regulating the cough reflex in guinea-pigs. J Physiol 2005; 569(Pt 2): 559–73.

57. Zhao Q, Yu CD, Wang R, Xu QJ, Dai Pra R, Zhang L, Chang RB. A multidimensional coding architecture of the vagal interoceptive system. Nature 2022; 603(7903): 878–84.

58. Cavanaugh DJ, Chesler AT, Braz JM, Shah NM, Julius D, Basbaum AI. Restriction of transient receptor potential vanilloid-1 to the peptidergic subset of primary afferent neurons follows its developmental downregulation in nonpeptidergic neurons. J Neurosci 2011; 31(28): 10119–27.

59. Stanford KR, Hadley SH, Barannikov I, Ajmo JM, Bahia PK, Taylor-Clark TE. Antimycin A-induced mitochondrial dysfunction activates vagal sensory neurons via ROS-dependent activation of TRPA1 and ROS-independent activation of TRPV1. Brain Res 2019; 1715: 94–105.

60. Lewis C, Neidhart S, Holy C, North RA, Buell G, Surprenant A. Coexpression of P2X2 and P2X3 receptor subunits can account for ATP-gated currents in sensory neurons. Nature 1995; 377(6548): 432–5.

61. Ellis JL, Undem BJ. Inhibition by capsazepine of resiniferatoxin- and capsaicin-induced contractions of guinea pig trachea. J Pharmacol Exp Ther 1994; 268(1): 85–9.

62. Mazzone SB, Geraghty DP. Respiratory actions of tachykinins in the nucleus of the solitary tract: characterization of receptors using selective agonists and antagonists. Br J Pharmacol 2000; 129(6): 1121–31.

63. Driessen AK, McGovern AE, Behrens R, Moe AAK, Farrell MJ, Mazzone SB. A role for neurokinin 1 receptor expressing neurons in the paratrigeminal nucleus in bradykinin-evoked cough in guinea-pigs. J Physiol 2020; 598(11): 2257–75.

64. Mazzone SB, Geraghty DP. Respiratory action of capsaicin microinjected into the nucleus of the solitary tract: involvement of vanilloid and tachykinin receptors. Br J Pharmacol 1999; 127(2): 473–81.

65. Ludwig MQ, Cheng W, Gordian D, Lee J, Paulsen SJ, Hansen SN, Egerod KL, Barkholt P, Rhodes CJ, Secher A, Knudsen LB, Pyke C, Myers MG, Jr., Pers TH. A genetic map of the mouse dorsal vagal complex and its role in obesity. Nat Metab 2021; 3(4): 530–45.

66. Lieu TM, Myers AC, Meeker S, Undem BJ. TRPV1 induction in airway vagal low-threshold mechanosensory neurons by allergen challenge and neurotrophic factors. Am J Physiol Lung Cell Mol Physiol 2012; 302(9): L941–8.

67. Watanabe N, Horie S, Michael GJ, Keir S, Spina D, Page CP, Priestley JV. Immunohistochemical co-localization of transient receptor potential vanilloid (TRPV)1 and sensory neuropeptides in the guinea-pig respiratory system. Neuroscience 2006; 141(3): 1533–43.

68. Watanabe N, Horie S, Michael GJ, Spina D, Page CP, Priestley JV. Immunohistochemical localization of vanilloid receptor subtype 1 (TRPV1) in the guinea pig respiratory system. Pulm Pharmacol Ther 2005; 18(3): 187–97.

69. Yamamoto Y, Sato Y, Taniguchi K. Distribution of TRPV1- and TRPV2-immunoreactive afferent nerve endings in rat trachea. J Anat 2007; 211(6): 775–83.

70. Hennel M, Harsanyiova J, Ru F, Zatko T, Brozmanova M, Trancikova A, Tatar M, Kollarik M. Structure of vagal afferent nerve terminal fibers in the mouse trachea. Respir Physiol Neurobiol 2018; 249: 35–46.

71. Farrell MJ, Bautista TG, Liang E, Azzollini D, Egan GF, Mazzone SB. Evidence for multiple bulbar and higher brain circuits processing sensory inputs from the respiratory system in humans. J Physiol 2020; 598(24): 5771–87.

72. Paton JF. Pattern of cardiorespiratory afferent convergence to solitary tract neurons driven by pulmonary vagal C-fiber stimulation in the mouse. J Neurophysiol 1998; 79(5): 2365–73.

73. Bonham AC, Joad JP. Neurones in commissural nucleus tractus solitarii required for full expression of the pulmonary C fibre reflex in rat. J Physiol 1991; 441: 95–112.

74. Vardhan A, Kachroo A, Sapru HN. Excitatory amino acid receptors in the nucleus tractus solitarius mediate the responses to the stimulation of cardio-pulmonary vagal afferent C fiber endings. Brain Res 1993; 618(1): 23–31.

75. Lovelace JW, Ma J, Yadav S, Chhabria K, Shen H, Pang Z, Qi T, Sehgal R, Zhang Y, Bali T, Vaissiere T, Tan S, Liu Y, Rumbaugh G, Ye L, Kleinfeld D, Stringer C, Augustine V. Vagal sensory neurons mediate the Bezold-Jarisch reflex and induce syncope. Nature 2023; 623(7986): 387-96.

76. Green JF, Schmidt ND, Schultz HD, Roberts AM, Coleridge HM, Coleridge JC. Pulmonary C-fibers evoke both apnea and tachypnea of pulmonary chemoreflex. J Appl Physiol Respir Environ Exerc Physiol 1984; 57(2): 562–7.

77. Schertel ER, Adams L, Schneider DA, Smith KS, Green JF. Rapid shallow breathing evoked by capsaicin from isolated pulmonary circulation. J Appl Physiol (1985) 1986; 61(3): 1237–40.

78. Raymond CS, Soriano P. High-efficiency FLP and PhiC31 site-specific recombination in mammalian cells. PLoS One 2007; 2(1): e162.

79. Kranz A, Fu J, Duerschke K, Weidlich S, Naumann R, Stewart AF, Anastassiadis K. An improved Flp deleter mouse in C57Bl/6 based on Flpo recombinase. Genesis 2010; 48(8): 512–20.

80. Zhao C, Ries C, Du Y, Zhang J, Sakimura K, Itoi K, Deussing JM. Differential CRH expression level determines efficiency of Cre- and Flp-dependent recombination. Frontiers in neuroscience 2023; 17: 1163462.

81. Choi S, Hachisuka J, Brett MA, Magee AR, Omori Y, Iqbal NU, Zhang D, DeLisle MM, Wolfson RL, Bai L, Santiago C, Gong S, Goulding M, Heintz N, Koerber HR, Ross SE, Ginty DD. Parallel ascending spinal pathways for affective touch and pain. Nature 2020; 587(7833): 258–63.

82. Patil MJ, Kim SH, Bahia PK, Nair SS, Darcey TS, Fiallo J, Zhu XX, Frisina RD, Hadley SH, Taylor-Clark TE. A Novel Flp Reporter Mouse Shows That TRPA1 Expression Is Largely Limited to Sensory Neuron Subsets. eNeuro 2023; 10(12).

83. Undem BJ, Taylor-Clark T. Mechanisms underlying the neuronal-based symptoms of allergy. J Allergy Clin Immunol 2014; 133(6): 1521–34.

84. Zaccone EJ, Undem BJ. Airway Vagal Neuroplasticity Associated with Respiratory Viral Infections. Lung 2016; 194(1): 25–9.

